# mRNA N1-2’O-methylation by CMTR1 affects NVL2 mRNA splicing

**DOI:** 10.1101/2023.09.28.560047

**Authors:** Steven Wolter, Thomas Hennig, Christina Wallerath, Thais M. Schlee-Guimaraes, Alexander Kirchhoff, Christian Urban, Antonio Piras, Alexey Stukalov, Stefan Juranek, Michael Engelke, Volker Boehm, Niels H. Gehring, Gunther Hartmann, Andreas Pichlmair, Caroline C. Friedel, Lars Dölken, Martin Schlee

## Abstract

Cap0-mRNA is characterized by a 5’-5’triphosphate-linked N7-methylated guanosine(m7G). In higher eukaryotes, the methyltransferase CMTR1 additionally methylates the 2’O-position of the penultimate mRNA nucleotide(N1) ribose (cap1-mRNA). While the m7G cap is essential for mRNA export and translation initiation by the eIF4F complex, the N1-2’O-methylation prevents recognition of cap1-mRNA by the antiviral RNA receptors RIG-I and IFIT1, but a function beyond immunotolerance remained elusive. Here, we generated *CMTR1*-knockout(*CMTR1*^-/-^) cells and found that type-I-interferon(IFN-I) treatment resulted in IFIT1-mediated reduction of cell viability and broad mRNA translation. Consequently, stimulation of the antiviral receptor RIG-I in *CMTR1*^-/-^ cells revealed an IFIT1 dependent dramatic reduction of IFN-I and chemokine protein induction, demonstrating the importance of N1-2’O-methylation for antiviral responses.

Additionally, IFN-I- and IFIT1-independent effects were observed: *CMTR1*^-/-^ cells were smaller, divided slower, and exhibited a reduced transcription of mRNAs coding ribosomal proteins (RP), 5’TOP-RNA and snoRNA host genes(SNHG). Additionally, proteome and transcriptome analysis revealed that expression of NVL2, an essential factor in ribosome biogenesis, is strongly suppressed by an alternative-splicing event of *NVL2 mRNA* in CMTR1^-/-^ cells. This reduction could only be rescued by catalytically active CMTR1. Altogether, besides antiviral immunity N1-2’O-methylation by CMTR1 has broad effects on cellular physiology and controls splicing of NVL2.

## INTRODUCTION

The mRNA cap plays a crucial role throughout the lifecycle of eukaryotic mRNAs: it stabilizes the mRNA against degradation and enables efficient mRNA maturation, export and translation (Jiao et al., 2013; Niedzwiecka et al., 2002; Ramanathan, Robb, et al., 2016; Sikorski et al., 2020).

Capping – the synthesis of an mRNA cap at the 5’end of the mRNA – is the earliest mRNA maturation step and occurs co-transcriptionally. In Metazoans, RNGTT transfers a Guanosin in 5’5 direction via a triphosphate linkage to the first nucleotide (N1) of the nascent pre-mRNA (GpppN1)(Ghosh et al., 2011; McCracken et al., 1997; Ramanathan, Robb, et al., 2016). RNMT methylates the Guanosin at position 7, generating the so called cap0 structure (m7GpppN1), (Aregger & Cowling, 2013; Ramanathan, Robb, et al., 2016; Tsukamoto et al., 1998). CMTR1 methylates the 2’O-position of the penultimate (N1) nucleotide, resulting in the cap1 structure (m7GpppN1m) (Belanger et al., 2010; Furuichi et al., 1975; Inesta-Vaquera & Cowling, 2017; Ramanathan, Robb, et al., 2016).

While cap1 is the dominant mRNA cap structure in mammals (Galloway et al., 2020; Wang et al., 2019), there are additional, less frequent cap methylations: CAPAM contranscriptionally methylates the N6 position of the penultimate Adenosin of certain mRNAs (Akichika et al., 2019; Boulias et al., 2019; Sendinc et al., 2019; Sun et al., 2019). Also, approximately 50% of mRNAs are posttranscriptionally methylated at the 2’O-position of the antepenultimate nucleotide(N2) by the cytoplasmatic methyltransferase CMTR2 (Furuichi et al., 1975; Werner et al., 2011).

Capping and transcription are tightly coupled: After transcribing the first 20-60 nucleotides, the RNA Pol II transcription complexes temporally halts (promoter proximal pausing) (Coppola et al., 1983; Core & Adelman, 2019; Rasmussen & Lis, 1993). Contemporaneously, capping enzymes are recruited to the C-Terminal domain (CTD) of RNA Polymerase II (RNA Pol II) and the mRNA cap is formed on the nascent RNA (Eick & Geyer, 2013; Kachaev et al., 2020). The mRNA cap is subsequently bound by the *cap binding complex* (CBC). Binding of CBC stimulates the transition of the transcription machinery into productive elongation and facilitates effective processing of the nascent RNA by protecting it against degradation as well as enhancing splicing, 3’end processing, export and the initiation of the first round of translation in the cytosol (Andersen et al., 2013; Flaherty et al., 1997; Jiao et al., 2013; Pabis et al., 2013; Ramanathan, Robb, et al., 2016). This complex interdependency of RNA polymerase II dependent transcription and capping is reviewed in detail elsewhere (Kachaev et al., 2020).

In the cytosol, the cap binding protein eIF4E binds to the mRNA cap and replaces CBC. eIF4E bound mRNA recruits further translation factors and is then efficiently translated into protein(Batool et al., 2019; Jackson et al., 2010).

Capping is an integral part of gene expression and highly conserved in eukaryotes. Interestingly, lower eukaryotes like yeast lack N1-2’O-methylation and therefore have cap0 modified mRNAs (Daugherty et al., 2016; Ramanathan, Robb, et al., 2016). While the mRNA cap is undoubtedly essential for the mRNA functionality, the effects mediated by CMTR1 catalysed N1-2’O-methylation are less clear: While both main CBC and eIF4E require m7G methylation (as found in cap0) for efficient binding, there does not seem to be a substantial preference for the additional N1-2’O-methylation in cap1 (Niedzwiecka et al., 2002; Tamarkin-Ben-Harush et al., 2017; Worch et al., 2005).

The decapping exonuclease DXO has been described to be involved in quality control of the mRNA cap: DXO degrades cap0, but not cap1 modified mRNA in vitro (Kramer & McLennan, 2019; Picard-Jean et al., 2018). The loss of DXO leads to an increase in incompletely processed mRNA in the cell (Jiao et al., 2013).

In early studies, N1-2’O-methylation has been described to enhance translation in different eukaryotic model systems (Kuge et al., 1998; Muthukrishnan et al., 1976, 1978). A more recent study suggests that differences in translational activity of cap0 and cap1 might be attributed to differences between the cell lines used(Sikorski et al., 2020).

The best described function of N1-2’O-methylation of mRNA is to mark host mRNA as endogenous in the context of antiviral immunity. Both the antiviral pattern recognition receptor RIG-I and the interferon inducible, antiviral effector have been shown to specifically bind to viral RNA with hypomethylated cap structures, including cap0, while tolerating endogenous, cap1 modified RNA(Bartok & Hartmann, 2020; Leung & Amarasinghe, 2016).

RIG-I efficiently gets activated by viral RNA in the cytosol and induces an antiviral type I Interferon (IFN-I) response. RIG-I detects double stranded RNA (dsRNA) ligands in a 5’dependent manner: 5’triphosphate (5’ppp), 5’diphosphat (5’pp) and cap0 modified RNAs activate RIG-I, while cap1 modified RNA does not elicit a response(Goubau et al., 2014; Devarkar, et al., 2016; Schlee et al., 2009; Schuberth-Wagner et al., 2015).

IFN-I induces the expression of *Interferon stimulated genes* (ISG) to combat invading viruses. IFIT1 is a highly IFN-I inducible, antiviral effector protein, that binds and restricts translation of single-stranded RNA (ssRNA) with hypomethylated mRNA caps like cap0, but not cap1 (Abbas et al., 2017; Leung & Amarasinghe, 2016; Pichlmair et al., 2011).

To elucidate the biological function of CMTR1 catalyzed N1-2’O-methylation of endogenous RNA we generated and characterized the CMTR1 knock-out (CMTR1^-/-^) in HEK293T cells. While the CMTR1^-/-^ cells are vital, they show reduced growth kinetic and smaller cell size.

IFN-I treatment caused an IFIT1-mediated strong reduction in viability and global translational inhibition. Following RIG-I stimulation with ppp-dsRNA, ISG protein production but not mRNA expression was strongly reduced by IFIT1 in CMTR1^-/-^ cells. These results demonstrate that the loss of N1-2’O-methylation leads to recognition of endogenous RNA species by IFIT1 and thereby would severely disrupt the effectiveness of an IFN-I dependent antiviral immune response.

Furthermore, we characterized the effect of N1-2’O-methylation of endogenous RNAs by transcriptome analysis. While no global effects of CMTR1 on transcript stability and splicing efficiency were detected, CMTR1^-/-^ cells showed a reduced transcript expression of components of the translational machinery. This was particularly striking for ribosomal protein (RP) encoding mRNA, but more broadly affected transcripts that harbour the conserved 5’terminal oligopryrimidine motif (5’TOP) and/or are snoRNA host genes (SNHGs). In addition, a highly specific CMTR1 dependent alternative splicing event of *NVL2* was identified. NVL2 has been described as an essential factor in ribosome biogenesis. In the absence of CMTR1, NVL2 pre-mRNA was alternatively spliced by a previously undescribed exon skipping event of the eighth exon, resulting in the formation of a non-functional isoform and strongly reduced NVL2 protein levels in CMTR1^-/-^.

Taken together, the results show that CMTR1 dependent N1-2’O-methylation is important to maintain self-tolerance of endogenous mRNA to IFIT-1, controls specific splicing events and affects the synthesis of components of the translation machinery on transcriptional level by an yet unknown mechanism.

## RESULTS

### CMTR1 deficient cells lack N1-2’O-methylation and show reduced cell proliferation and cell size

The majority of capped transcripts are methylated at both the 7G and N1-2’O-position, thereby making cap1 (Fig. 1A)) the dominant mRNA cap structure in vertebrate cells(Galloway et al., 2020; Wang et al., 2019).

**Figure 1:**
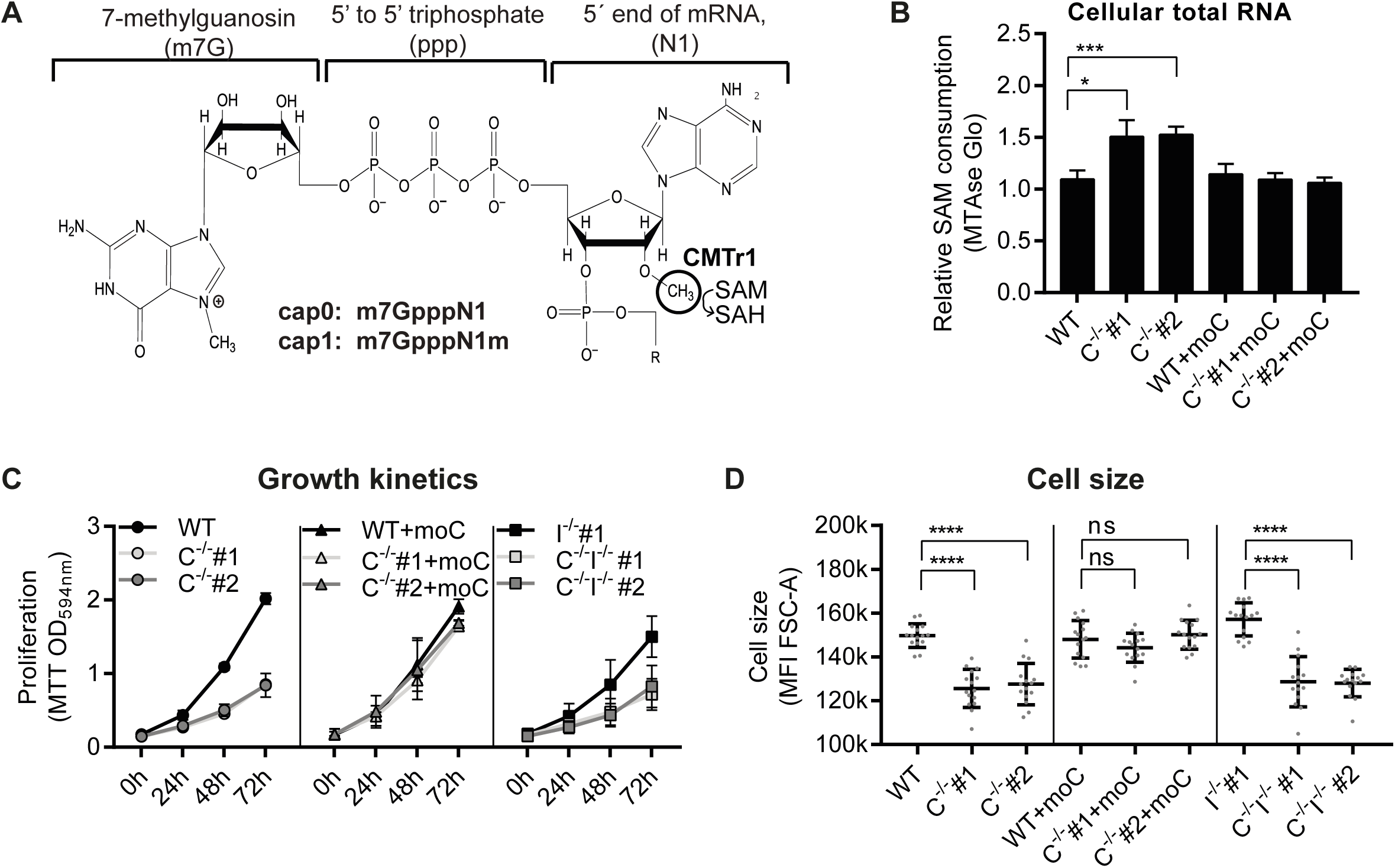
CMTR1^-/-^ cells lack N1-2’0-methylation and have reduced growth kinetic and cell size. A) Structure of the mRNA Cap. Shown are the 7-methyl guanosin cap (m7G) and the first two nucleotides (N1 and N2) of an mRNA. The methylation of the N1-2’0 position by CMTr1 is indicated. B) 0etection of endogenous capO-modified RNA with the MTAseGlo Assay. The relative SAM consumption is measured as the quotient of luminescence intensities between enzyme and mock reaction. 2OOOng RNA of cellular total RNA were used (n=4, mean +SD). C) Analysis of cell growth kinetics. 5×10^5^ cells were plated and metabolic activity was measured by MTT at the indicated timepoints for 1h. MTT turnover was measured photospectrometrically at 00594nm. (n=5, mean +SD) 0) Flow cytometric analysis of cell size. The mean fuorescence intensity of FSC-A of the cell population is plotted (n=16, mean + SD)

To study the effect of N1-2’O-methylation we generated CMTR1 deficient HEK293T cell lines (*C*^-/-^ *#1* and *C*^-/-^ *#2*) by CRISPR/Cas9 knock-out. As a control, we stably expressed murine *CMTR1* (*moC*) from a retroviral integrated expression cassette in HEK293T wildtype cells (WT), C^-/-^#1 and C^-/-^ #2, resulting in the cell lines *WT+moC*, *C*^-/-^ *#1+moC and C*^-/-^ *#2+moC*, respectively. To confirm loss of N1-2’O-methylation after *CMTR1* knock-out (Belanger et al., 2010; Lee et al., 2020) we established a non-radioactive assay (MTAseGlo) that asses the N1-2’O-methylation status of RNA (Fig.1B and Suppl. Fig. 1A-C).

The lack of N1-2’O-methylation in CMTR1^-/-^ cells makes endogenous mRNA to bona fide ligands for the antiviral effector protein IFIT1, that is known to bind and sequester RNA with hypomethylated cap structures from translation (Abbas et al., 2017; Habjan et al., 2013; Leung & Amarasinghe, 2016). While IFIT1 has a low basal expression, IFIT1 is highly induced upon IFN-I treatment. To study the effect of IFIT1 on the phenotype of CMTR1^-/-^, we knocked out IFIT1 in WT, C^-/-^#1 and C^-/-^ #2, resulting in the IFIT1 deficient celline (IFIT1^-/-^) I^-/-^, and the CMTR1 and IFIT1 deficient cellines (CMTR1^-/-^IFIT1^-/-^) C^-/-^I^-/-^ #1 and C^-/-^ I^-/-^ #2, respectively.

First, we analyzed the growth kinetics of CMTR1^-/-^ cells. We measured the metabolic activity as a surrogate parameter for total cell number on three consecutive days after seeding and found that CMTR1 deficient cells, independently of the IFIT1 status, have a significantly reduced growth kinetic (Fig. 1C).

FACS analysis revealed that CMTR1 deficient cells, independently of the IFIT1 status, were significantly reduced in cell size, as indicated by a shift of the population in the FSC-A (Fig. 1D). While it is known that the cell size changes during cell cycle progression (Björklund, 2019; Lloyd, 2013), the difference in cell size between the tested cell lines was not caused by changes in the cell cycle profile (Supplementary Fig. 1D+E).

### IFN-α treatment results in IFIT1 mediated reduced viability and translational inhibition in CMTR1**^-/-^** cells

IFN treatment puts the cells in an antiviral state by inducing the expression of IFN stimulated genes (ISGs), including antiviral pattern recognition receptors like RIG-I and antiviral effector proteins like IFIT1(Fensterl & Sen, 2015; Ivashkiv & Donlin, 2014). To monitor metabolic activity and thus viability of cells we performed the colorimetric MTT assay 7 days after IFN-α treatment and found a significant IFN dose-dependent reduction of viability in CMTR1^-/-^ cells (Fig. 2A, C^-/-^). This effect was completely compensated in cells deficient for both CMTR1 and IFIT1 (Fig. 2A, C^-/-^I^-/-^).

**Figure 2:**
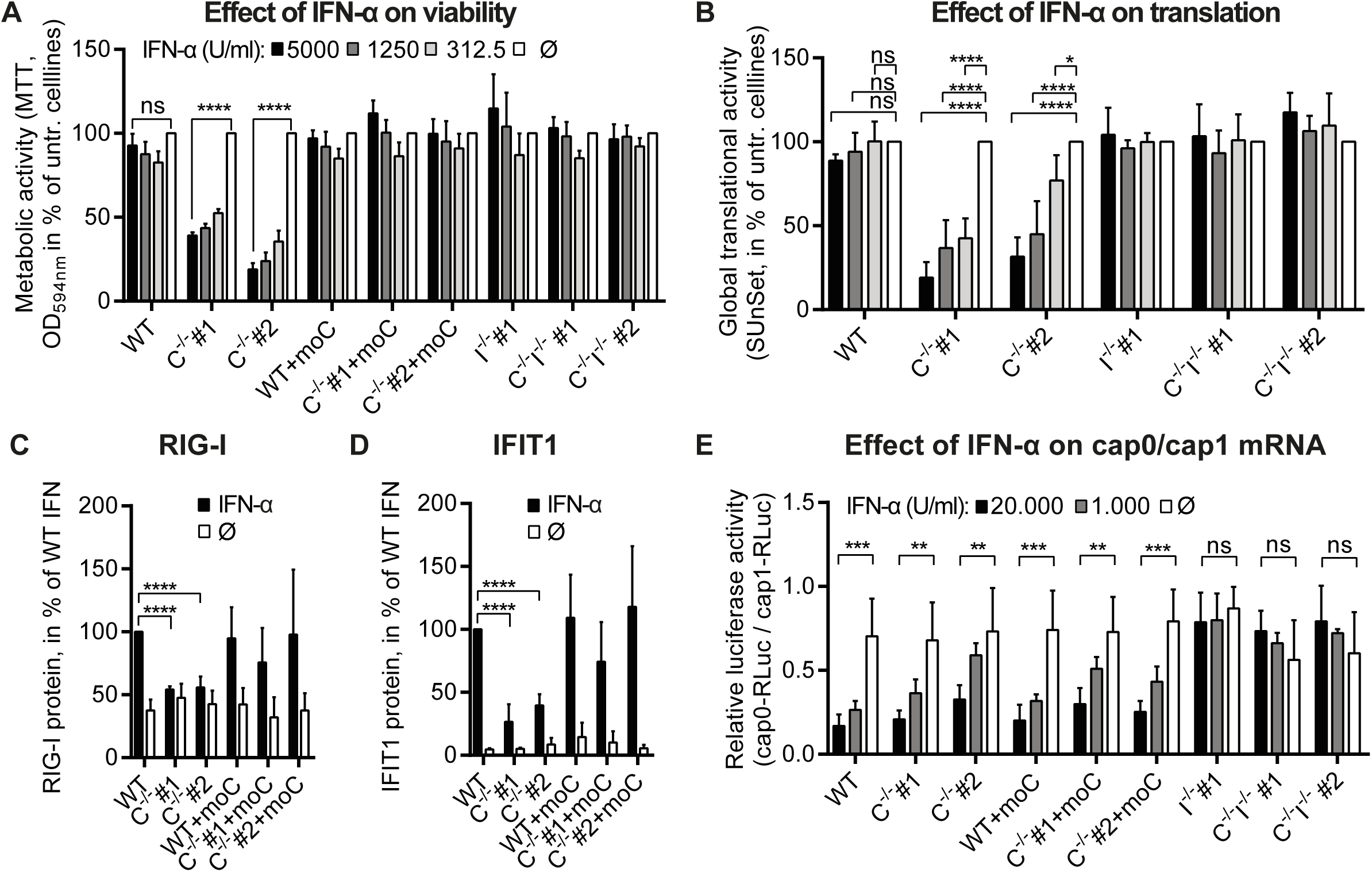
IFIT1 mediated Effects in IFN-α treated CMTR1^-/-^ cells. A) 5×10^3^ cells/well (96 well plate) were seeded and treated with the indicated IFN-α concentration After 7d the metabolic activity was measured using MTT Assay. The results are normalized to the untreated control of each cell line (n=6, mean + SD) B) Global Translational Activity 2×10^5^ cells were plated in 6 well plates and treated with the indicated IFN-α concentrations After 72h, SUNSet assay was performed The signal intensity of the puromycin western blot (WB) was normalized to the signal intensity of the Ponceau staining for each lane The results are shown in % of the untreated control of each cell line C+D) 2*10^5^ cells were plated in 6 well plates and treated with the 5OOOU/ml IFN-α or left untreated After 72h, cells were lysed and analyzed by western blot (WB) The signal intensity of the RIG-I WB (C) and IFIT1 WB (0) were normalized to the Actin WB Results in % of untreated WT cells E) Effect of N1-2’0-methylation on translation of reporter mRNA 2*10^5^ cells/well (96 well plate) were treated with indicated concentrations of IFN-α for 24h, then transfected with 2Ong capO or cap1-modified RLuc reporter mRNA After 24h, luciferase activity was measured The relative luciferase activity is calculated as the ratio of capO-RLuc to cap1-RLuc luciferase activity (n=4, mean +SD)

We confirmed this effect of IFN-α treatment by a *colony formation assay* (CFA) (Suppl. Fig. 2). CFA allows estimation of colony number and size. Here, the effect of CMTR1 k.o. and the additional effect by IFN-I treatment became visible: In untreated condition, the size of individual colonies of CMTR1^-/-^ cells (*C*^-/-^ *#1/#2)* and CMTR1^-/-^IFIT1^-/-^ (*C*^-/-^ *I*^-/-^ *#1/#2)* cells were equally reduced compared to CMTR1-expressing cells (WT, *WT+moC*, *C*^-/-^ *#1+moC, C*^-/-^ *#2+moC*). This is in agreement with the observed reduced growth kinetics (metabolic activity) of CMTR1 deficient cells (Fig. 1C). Importantly, IFN-α treatment further reduced the growth of CMTR1^-/-^ cells. In contrast, the growth of CMTR1^-/-^IFIT1^-/-^ and CMTR1-expressing cells was, if at all, only mildly affected by IFN-α treatment.

Since IFIT1 restricts the translation of bound cap0 modified RNA (Habjan et al., 2013), we next determined the effect of IFN-α on the global translation activity of the cells by *SUnSet* (Surface Sensing of Translation) assay that monitors puromycin incorporation in actively translated proteins by western blot. The translational activity of CMTR1^-/-^ was significantly reduced by IFN-α treatment for 72h in a dose-dependent manner, while WT as well as CMTR1^-/-^IFIT1^-/-^ cells were unaffected by IFN-α treatment (Fig. 2B). Accordingly, the IFN-a mediated induction of the ISGs IFIT1 and RIG-I on protein level was strongly impaired (Fig. 2C/D).

To analyze effects of N1-2’O-methylation on RNA translation separated from RNA transcription and processing, the translation of exogenous cap0 and cap1 reporter mRNA was compared. Cells were pre-treated with IFN-α or control and then transfected with cap0- or cap1-modified Renilla luciferase (RLuc) mRNA. 24 h after transfection, the Luciferase activity was determined.

IFN-α treatment strongly decreased the relative cap0 reporter luciferase activity (cap0/cap1) in all IFIT1-competent cells but not in IFIT1-deficient cells (Fig. 2E). Of note, this effect was not dependent on the CMTR1 status of the cell. Remarkably, in the absence of IFN-α, the relative cap0/cap1 luciferase activity was equal among all genotypes (mean relative cap/cap1 luciferase activity across all cell lines 70% (+/- 9% SD)).

Altogether, This demonstrates that the used CMTR1^-/-^ cell lines have not acquired compensatory genetic alterations that enable or enhance translation of cap0-RNA.

### IFIT1 blocks the translation of RIG-I induced IFN-I and the chemokine CXCL10 in CMTR1 deficient cells

Apart from IFIT1, the antiviral receptor RIG-I is central for in innate immunity against RNA viruses. Activation of the RIG-I signaling pathway results in the production of chemokines such as CXCL10 and IFN-I that subsequently induces the expression of antiviral genes (Bartok & Hartmann, 2020). We stimulated the cells with exogenic RIG-I ligand to analyze whether the RIG-I pathway is altered in CMTR1 deficient cells. After stimulation with ppp-dsRNA (IVT4), we assessed the RIG-I response by measuring mRNA expression and secretion (reporter bioassay) of IFN-I and CXCL10 (ELISA). Strong induction of IFN-beta as well as CXCL10 mRNA (Fig. 3A, B upper panels) and phosphorylation of TBK1 (Fig. 3D) in all cell lines confirmed that RIG-I signaling and gene expression induction were not impaired in CMTR1^-/-^ cells. However, RIG-I mediated CXCL10 secretion and IFN-I bioactivity were dramatically reduced in CMTR1^-/-^ cells (Fig. 3A,B lower panels). Intriguingly, this effect was almost compensated in CMTR1^-/-^IFIT1^-/-^ cells (Fig. 3A,B lower panels). Of note, since CMTR1 deficiency causes also independently of IFN-I slower proliferation in CMTR1^-/-^IFIT1^-/-^ (see Fig. 1C), CXCL10 and IFN-I concentration are reduced due to reduced absolute cell numbers per well, thereby underestimating the RIG-I activation in CMTR1^-/-^IFIT1^-/-^ cells (Fig. 3A/B). Similar to CXCL10 and IFN-I, the induction of protein but not mRNA of IFIT1 and RIG-I were significantly reduced in CMTR1^-/-^ cells (Fig. 3B, Suppl. Fig. 3A-C).

**Figure 3:**
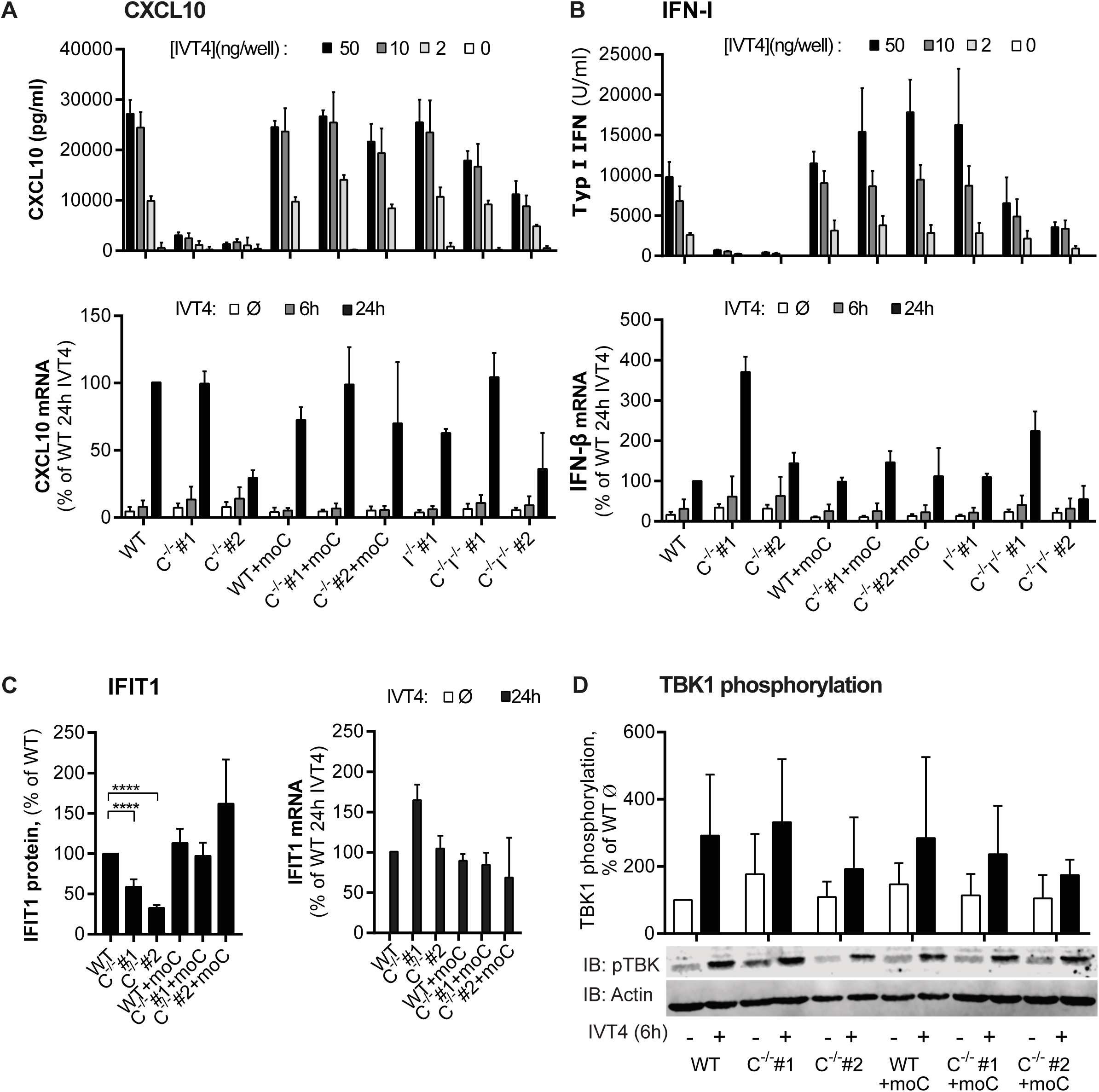
CMTR1^-/-^ cells show reduced protein production after RIG-I stimulation. A)/B) Upper panels: Cells were transfected with the indicated amount of IVT4. After 24h, the concentration of IP10 (ELISA) and activity of IFN-I (Hekblue IFN reporter) in supernatants were measured. Results in % to WT 50ng IVT4 (n=3-4, mean +SD). A)/B) lower panels: mRNA expression qPCR analysis of CXCL10 and IFN-β in indicated cells stimulated with IVT4 (500 ng/ml) for 6h or 24h or left untreated. Expression normalized to GAPDH is shown (n=4, mean + SD). C) Left panel : 2×10^5^ indicated cells were stimulated with IVT4 (1ug/ml). After 24h, protein levels of IFIT1, normalized to Actin, were determined by western blot and quantified. Results in % of WT (n=3, mean +SD). Right panel : mRNA expression qPCR analysis of IFIT1 in indicated cells stimulated with IVT4 (0.5 ug/ml) for 24h or left untreated. Expression normalized to GAPDH is shown (n=4, mean + SD). D) Analysis of TBK1 phosphorylation after RIG-I stimulation: 6h after IVT4 (1 µg/ml) stimulation the phosphorylation of TBK1 (pTBK1) was quantlified by western blot. One representative western blot of pTBK1 and Actin is shown. Quantification of the relative amount of phosphorylated TBK1 (pTBK1) normalized to actin (n=6), mean + SEM.

Our results suggest that translational inhibition by IFIT1 would prevent the production of cytokines and antiviral effector proteins downstream of RIG-I or other PRR on the translational level if mRNA was not N1-2’O-methylated. Consequently, effective IFN-I dependent antiviral immunity is expected to be compromised in CMTR1 deficient cells. In agreement with this, siRNA mediated knockdown of CMTR1 has been shown to enhance viral replication of Zika and Vesicular Stomatitis Virus (Williams et al., 2020)

### CMTR1 shows no global effect on splicing and transcript stability but affects transcription of ribosomal proteins

To gain further insight into the phenotype of CMTR1^-/-^ cells, we performed genome wide RNA-sequencing (RNA seq). In general, defects in mRNA capping are expected to impact further maturation of mRNA (Kachaev et al., 2020). To assess possible effects of CMTR1 on splicing and RNA stability, we included both total cellular RNA (total RNA) and newly synthesized RNA Samples in the experiment. Newly transcribed cellular RNA was isolated by metabolic labeling with incorporating 4-thiouridine (4sU) for 1h before RNA extraction and subsequent thiol specific affinity purification as described before (Fig. 4A) (Dölken et al., 2008). The newly transcribed RNA is hereinafter referred to as *4sU RNA*.

**Figure 4:**
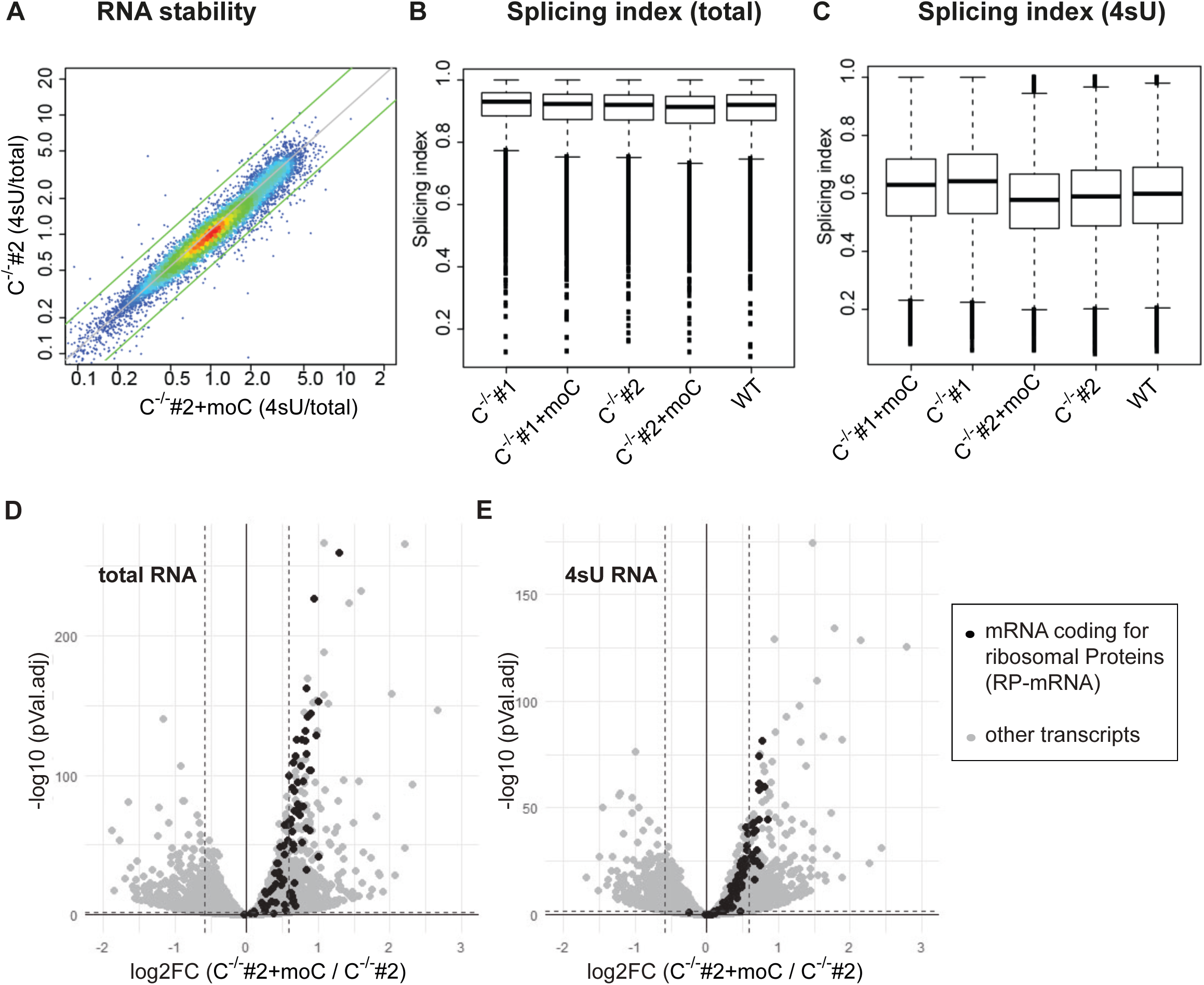
Analysis of the effects of CMTR1 on RNA stability, splicing and differential gene expression by transcriptome analysis. RNA-seq of total RNA (total) and newly synthesized, 4sU containig RNA (4sU)l. A) Analysis of the effects of CMTr1 on global transcript stability (4su/total). Representative result of the comparison of transcript stability in C^-/-^#2+moC vs C^-/-^#2l. C,D) Analysis of the global splicing eficiency (splicing index) for total RNA (B) and 4sU-RNA (C)l. D+E) Differential transcript expression between C^-/-^#2+moC and C^-/-^#2 in total RNA (D) and 4sU RNA (E)l. RP transcripts are shown as black dots, all other transcripts as grey dotsl. The dashed auxiliary lines mark log2FC = -0l.586 and 0l.586 and -log10 (pVal.adj) = 1l.311 (corresponding to pVal.adj of 0l.05)l.

The stability of a given RNA is dependent on its synthesis and degradation rates. We used the quotient of transcript expression values in 4sU RNA and versus total RNA (4sU/total) to calculate the relative stability of all measured transcripts in each cell line. Next, we plotted the relative stability of transcripts in CMTR1^-/-^ versus CMTR1^+/+^ cells but could not find a global effect of CMTR1 on RNA stability (Fig. 4A).

To assess possible effects of CMTR1 on splicing, we calculated the splicing index for each cell line by using the sequence information of reads that overlap with annotated splicing sites and divided exon-exon spanning reads -indicating a completed splicing reaction-by the total number of reads spanning splicing sites. As expected, the splicing index was high in total RNA and lower in 4sU RNA. However, we could not detect differences between CMTR1 deficient and CMTR1 competent cell lines (Fig. 4B+C).

We next determined the differentially expressed transcripts between CMTR1-competent and CMTR1^-/-^ cell lines. For this purpose, the conditions listed in Table 2 were compared.

**Table 1.**
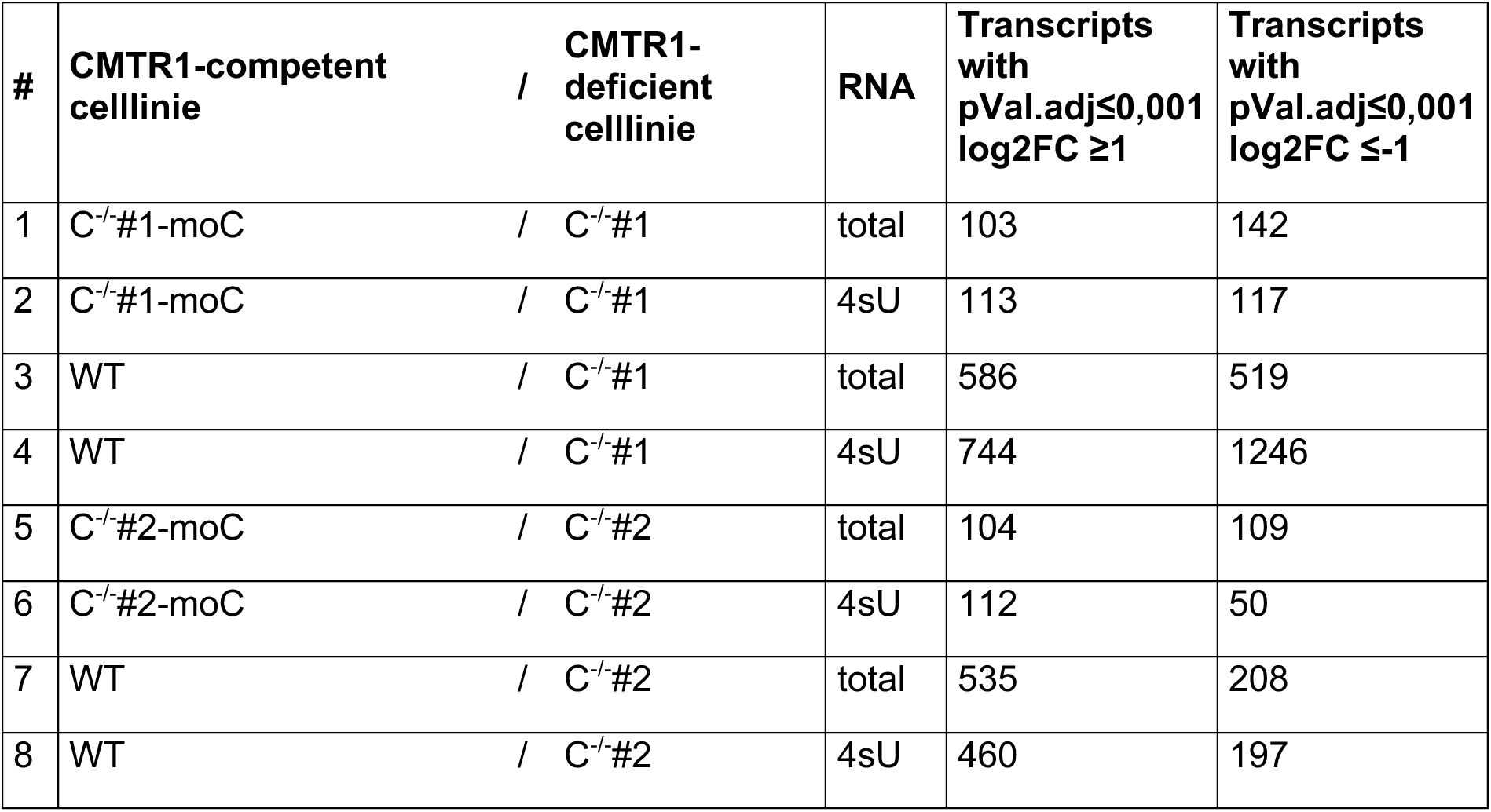
Comparison of indicated cell lines to determine differentially expressed transcripts. A total of 8 comparisons of transcript expression were made between the indicated CMTR1-competent and CMTR1-deficient cell lines. The number of transcripts significantly decreased expressed in CMTR1 (pVal.adj<0.001 log2FC >1) and significantly increased expressed in CMTR1 (pVal.adj<0.001 log2FC <-1) is indicated

| # | CMTR1-competent cellline | / CMTR1-deficient cellline | RNA | Transcripts with pVal.adj≤0,001 log2FC ≥1 | Transcripts with pVal.adj≤0,001 log2FC ≤-1 |
| --- | --- | --- | --- | --- | --- |
| 1 | C<sup>-/-</sup>#1-moC | / C<sup>-/-</sup>#1 | total | 103 | 142 |
| 2 | C<sup>-/-</sup>#1-moC | / C<sup>-/-</sup>#1 | 4sU | 113 | 117 |
| 3 | WT | / C<sup>-/-</sup>#1 | total | 586 | 519 |
| 4 | WT | / C<sup>-/-</sup>#1 | 4sU | 744 | 1246 |
| 5 | C<sup>-/-</sup>#2-moC | / C<sup>-/-</sup>#2 | total | 104 | 109 |
| 6 | C<sup>-/-</sup>#2-moC | / C<sup>-/-</sup>#2 | 4sU | 112 | 50 |
| 7 | WT | / C<sup>-/-</sup>#2 | total | 535 | 208 |
| 8 | WT | / C<sup>-/-</sup>#2 | 4sU | 460 | 197 |

**Table 2:** GO Enrichment results for "cytosolic ribosome". GO enrichment is based on classification of genes by Protein Analysis Through Evolutionary Relationships (PANTHER). Single GO Enrichments for differentially regulated genes (criteria log2(FC) >0.586 or <-0.586 ; pVal.adj <0.05) between the indicated cell lines (respectively CMTR1-competent cell line/CMTR1-deficient cell line) were performed and the results of the gene family "cytosolic ribosome" (GO term 0022626) were listed. Indicated are x-fold overrepresentation (fold enrichment), direction of regulation (overrepresentation/underrepresentation (+/-)), and significance (pVal (p-value) and FDR (false discovery rate)). The analysis was performed using the PANTHER web application (http://www.pantherdb.org).

| CMTR1-competent celllinie / CMTR1-deficient celllinie | RNA | Results of GO Enrichment für "cytosolic ribosome" (GO term GO:0022626) |  |  |  |
| --- | --- | --- | --- | --- | --- |
|  |  | fold enrichment | +/- | pVal | FDR |
| C<sup>-/-</sup>#1-moC / C<sup>-/-</sup>#1 | total | 20,86 | + | 1,86E-36 | 3,74E-33 |
|  | 4sU | 6,25 | + | 4,85E-07 | 0,000974 |
| WT / C<sup>-/-</sup>#1 | total | 7,27 | + | 8,19E-24 | 1,65E-20 |
|  | 4sU | 3,77 | + | 7,31E-09 | 2,94E-06 |
| C<sup>-/-</sup>#2-moC / C<sup>-/-</sup>#2 | total | 21,71 | + | 6,65E-43 | 1,33E-39 |
|  | 4sU | 6,75 | + | 9,79E-09 | 1,97E-05 |
| WT / C<sup>-/-</sup>#2 | total | 7,61 | + | 7,63E-26 | 1,53E-22 |
|  | 4sU | 6,92 | + | 8,69E-19 | 1,75E-15 |

We used GO enrichment to identify differentially regulated classes of genes and found a highly significant and almost uniformly downregulation of the GO term “cytosolic ribosome” (Table 1, Fig. 4D,E) in CMTR1^-/-^ cells. This GO term comprises mostly structural components of the ribosome (Ribosomal proteins (RP)). Other significantly regulated GO terms strongly overlap with the GO term “cytosolic ribosome” and are not listed here.

These results were unexpected, since RP are known to be mostly regulated on translational level, while RP mRNA is mostly constitutively expressed and individual RP mRNAs are routinely used as a housekeeping gene in qPCR experiments (Meyuhas & Kahan, 2015).

Next, we aimed at characterizing the mechanisms of RP mRNA downregulation in CMTR1^-/-^. Compared to other genes, RP-encoding genes have two well-characterized peculiarities: First, RP genes encode so-called 5’terminal oliopyrimidine (5’TOP) transcripts. Second, many RP genes, as well as other representatives of 5’TOP RNA coding genes, are snoRNA host genes (SNHGs), i.e. encode one or more snoRNAs within their intron sequences (Meyuhas & Kahan, 2015). During splicing of the SNGS, snoRNAs are processed from the intron sequence and are maturated. SnoRNAs guide the 2’O-methylation and pseudouridylation of specific residues within the RNA component of the ribosome (rRNA), thereby contributing to biosynthesis and function of the ribosome (Kufel & Grzechnik, 2019; J. Liang et al., 2019; Ojha et al., 2020).

We now assessed whether RP- and non-RP transcripts featuring 5’TOP and SNHG elements are differentially expressed in a CMTR1-dependent manner. For this purpose, the corresponding transcripts were annotated using published data (see below).

5’TOP Transcripts are defined structurally by the 5’TOP motif (the first nucleotide of the mRNA is a C followed by 4-15 pyrimidines (C/U)), and functionally by its translational regulation through the mTOR pathway (Reference!). Approximately 100 genes meet both the structural and functional definition of 5’TOP RNA ("validated 5’TOP RNA"), including mainly proteins of the translational machinery: 79 RPs as well as various initiation factors (EIF3A, EIF3E, EIF3H, EIF4B, PABPC1) and elongation factors (EEF1A1, EEF1B2, EEF1D, EEF1G, and EEF2)). The validated 5’TOP genes were annotated and divided into ribosomal 5’TOP (RP) and non-ribosomal 5’TOP (non-RP) (Meyuhas & Kahan, 2015; Philippe et al., 2020).

In addition to established 5’TOP genes, other genes whose transcripts harbor the 5’TOP or related motifs but haven’t been demonstrated to be translationally regulated by mTOR were defined as *5’TOP-like* genes (Philippe et al., 2020).

SNHGs are both protein-coding and non-coding genes in whose transcripts one or more snoRNA are localized. snoRNA is maturated from the intron sequence after splicing of the SNHG transcript (Kufel & Grzechnik, 2019). 5’TOP genes are above average frequency SNHGs. SNHGs were annotated according to the snoDB database (Bouchard-Bourelle et al., 2020).

In addition to protein-coding RP genes, there are over 2000 noncoding but partially transcribed ribosomal pseudogenes (pseudo-RPs) in humans. Pseudo-RPs have also been annotated (Zhang et al., 2002).

Based on the criteria described by (Philippe et al., 2020), genes were hierarchically classified into non-overlapping groups:

1) RP: protein-coding genes encoding ribosomal proteins.
2) non-RP 5’TOP: protein-coding genes with validated 5’TOP encoding non-ribosomal proteins.
3) ncSNHG 5’TOP-like: non-coding SNHGs with TOP score >3
4) ncSNHG non-5’TOP: non-coding SNHGs with TOP score <3
5) 5’TOP-like: coding and non-coding genes with a TOP score >3 that are not already included in group 1-5.
60 pseudo-RP: non-coding ribosomal pseudogenes not already included in group 1-6.
7) snoRNA: transcripts of the biotype "snoRNA".

Differential expression of these transcript groups was visualized using the C^-/-^#1+moC /C^-/-^#1 comparison pair as an example (Fig. 5A-H). All groups showed decreased expression in CMTR1-deficient cells. Comparison of ncSNHG-5’TOP-like and ncSHNG non-5’TOP suggests an additive effect of 5’TOP motif and SNHG status on downregulation of a transcript in CMTR1-deficient cells (Fig. 5C,D). Interestingly, transcripts of RP pseudogenes are also downregulated (Fig. 5F). Expression levels of snoRNAs are also reduced in CMTR1-deficient cells (Fig. 5G). This does not necessarily result from reduced SNHG expression, because snoRNA expression levels were found to correlate only conditionally with the corresponding SNHGs (Kufel & Grzechnik, 2019). To validate the results, the relative expression levels of selected target genes were analyzed by qPCR. In RNA extracted from untreated cell lines, the relative transcript expression of ribosomal proteins (RPS12, RPL13A, RPL10, RPS8, RPS23, RPLP10), elongation factors (EEF1A), noncoding SNHGs (SNHG8, SNHG19, SNHG12), snoRNAs (SNORA24, SNORD99), and phospholipase C Eta 1 (PLCH1) were found to be significantly reduced in CMTR1^-/-^cells compared with WT cells (Suppl. Fig. 4). We therefore conclude that deficiency of CMTR1 affects not only RP mRNA, but also other transcripts involved in the translational machinery of the cell.

**Figure 5:**
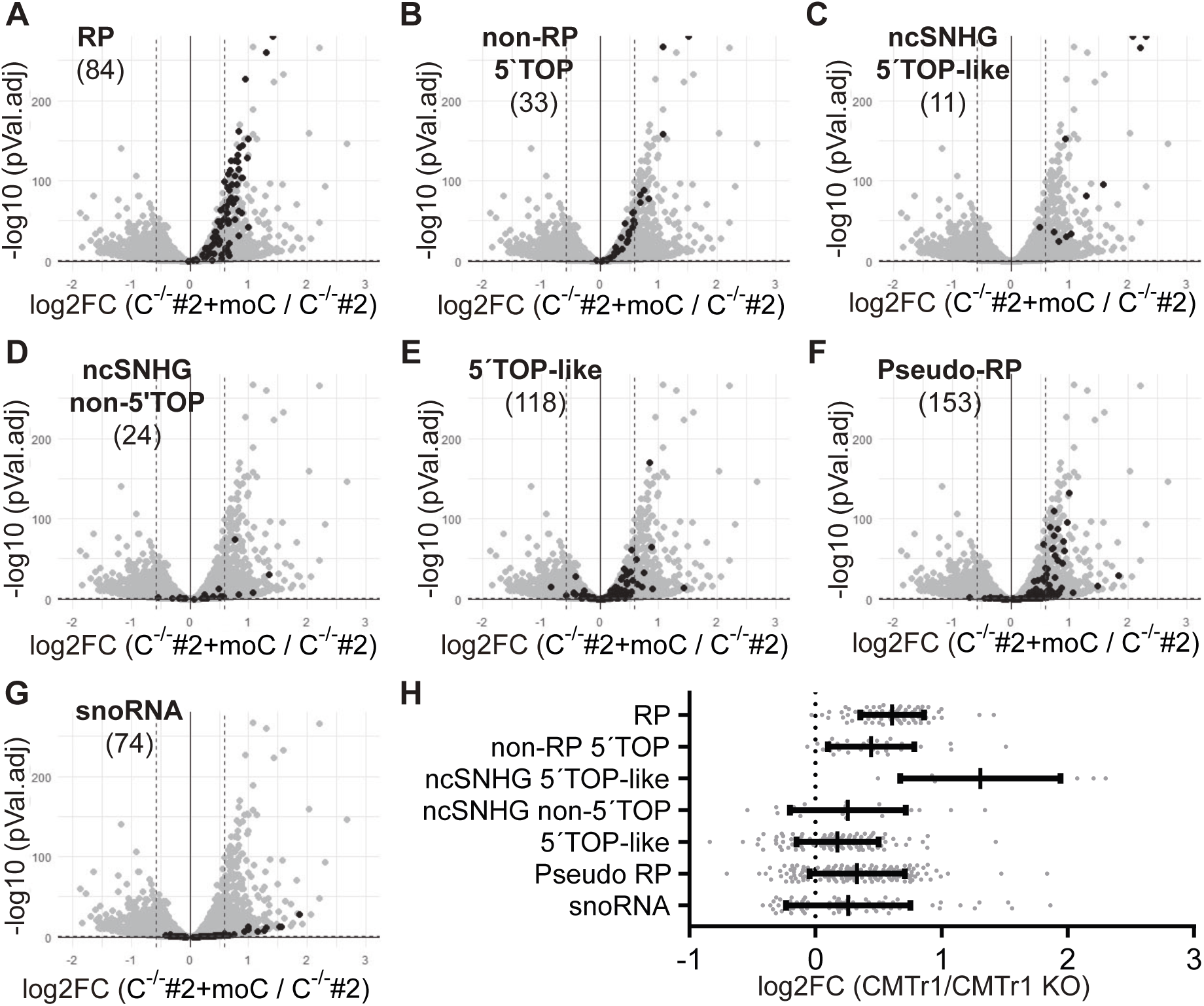
Transcripts coding ribosomal proteins, SNHGs, 5’TOP transcripts and snoRNAs are reduced in CMTR1^-/-^ cells. Differentia transcript expression between C^-/-^#2+moC and C^-/-^#2 in tota RNA. Transcripts were ordered and ana yzed in non-over apping groups based on pub ished datal. The groups are ribosoma protein (RP) mRNA (A), non-RP mRNA with 5’TOP motif (non-RP 5’TOP) (8), non-coding SNHG with TOP score >3 (ncSNHG 5’TOP) (C) and with TOP score <3 (ncSNHG non-TOP) (D), other transcripts with TOP score >3 (5’TOP ike) (E), transcripts of ribosoma pseudogenes (pseudo-RP) (F) and snoRNA (snoRNA) (G)l. Transcripts of the individua group are shown as b ack dots, a other transcripts as grey dotsl. The number of transcripts in the group is given in brackets be ow the group namel. The auxi iary ines mark og2FC= -0l.586 and 0l.586 as we as - og10(pVa l.adj)= 1l.311 (corresponds to pVa l.adj of 0l.05)l. H) P ot of og2FC of the groups as a scatter dot p ot (median with interquarti e range)l.

### CMTR1 deficiency causes alternative splicing with a premature termination codon in NVL2 mRNA

We next performed a proteome analysis of CMTR1-deficient (C^-/-^#1) and CMTR1-competent (C^-/-^#1+moC) cells. Of the total 7590 proteins identified and quantified, 48 proteins were differentially (pVal.adj <0.05. Log2FC >1 or <-1) regulated. No RPs were among the significantly differentially expressed proteins, however, the visualization of RPs within all detected proteins suggests a small but uniform reduction of RP expression in CMTR1^-/-^ cells (Fig. 6A).

**Figure 6:**
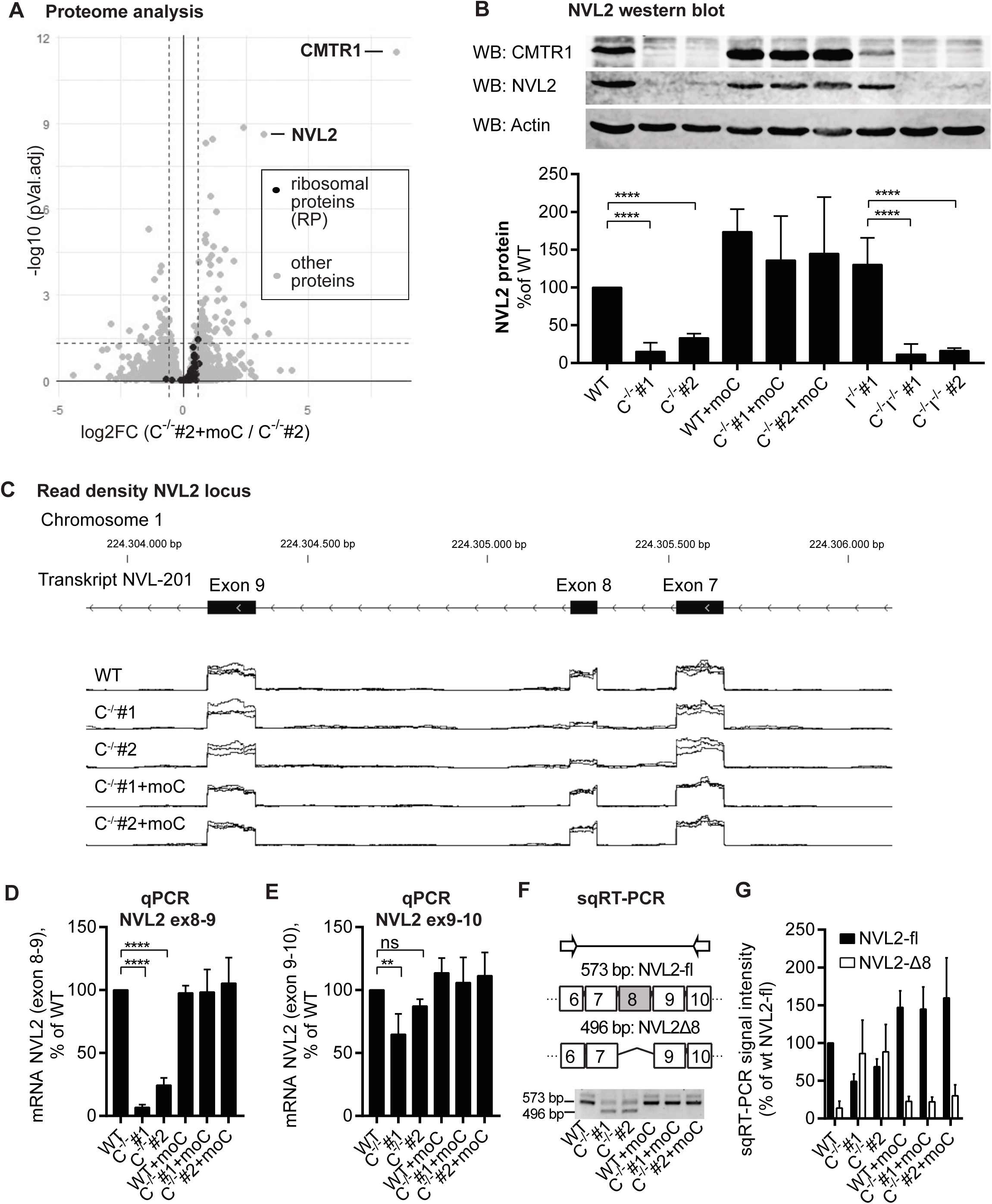
Reduced NVL2 expression in CMTR1^-/-^ cells is due to an alternative splicing event. A) Analysis of the differential protein expression between C^-/-^#2+moC and C^-/-^#2 cells by mass spectrometry (LFQ)l. Ribosomal proteins (RP) are shown as black dots, all other transcripts as grey dotsl. The data points of the proteins CMTr1 and NVL2 are markedl. The dashed auxiliary lines mark log2FC = -0l.586 and 0l.586 and - log10(pVal.adj)=1l.311 (corresponding to pVal.adj of 0l.05)l. B) Protein levels of CMTR1 and NVL2, normalized to Actin, were determined by western blotl. (Representative result, n=4, mean +SD)l. C) Representation of the read-density of a segment of the NVL2 gene locus (chromosome 1, 224,304,000-224,306,000, annotated with transcript NVL-201 exon 7-9) based on the RNA-seql. Superimposed read-density of all replicates per cell line (total RNA) D-F) Detection of NVL2 exon 8 exon skipping in RNA of untreated cellsl. Determination of relative transcript levels of NVL2 by exon 8 skipping sensitive (D) and insensitive (E) qPCR, normalized to GAPDH, in % of WT (n=4, mean + SEM)l. G) Representative experiment and evaluation of sqRT-PCRl. Shown is the relative intensity of the bands in % of wt (n=4, mean + SEM)l.

In CMTR1^-/-^ cells, besides CMTR1 it-self we observed the strongest reduction for *Nuclear Valosin-Containing Protein-Like* 2 (NVL2). Since NVL2 was described to play an important role in ribosome biogenesis (Hiraishi et al., 2018; LaCava et al., 2005; Yoshikatsu et al., 2015) it represents a putative candidate for CMTR1-mediated effects on transcript levels of RPs, SNHGs, and snoRNA seen in the RNA-seq dataset.

Western blot confirmed that NVL2 was strongly reduced in CMTR1-deficient cells (Fig. 6B). Subsequently, the expression of NVL2 at the transcript level was analyzed in the RNAseq data: In contrast to a strong reduction of NVL2 protein, NVL2 mRNA within total RNA showed only a moderate reduction in CMTR1-deficient cells and was even less altered in 4sU RNA. The difference between total and 4sU RNA expression levels of NVL2 indicates a reduced half-life of NVL2 transcripts in CMTR1-deficient cells.

The NVL2 gene locus indicates the cause of the altered transcript levels and stability of NVL2: Two protein-coding isoforms of NVL2 have been described so far (Nagahama et al., 2004): The dominant 858 amino acids (AS) NVL2 isoform is encoded by the NVL-201 transcript which comprises 23 exons and corresponds to the protein size detected in the western blot. The second described protein-coding transcript NVL-203 lacks exon 2 and encodes a 750 AS NVL2 isoform. Analysis of read density across the genomic locus of NVL2 revealed no CMTR1-dependent change in exon 2 expression (data not shown), but a strong reduction of exon 8 in CMTR1-deficient cells compared with CMTR1-competent cells in both total RNA and 4sU RNA (Fig. 6C).

An NVL2 transcript with exon skipping of exon 8 - hereafter referred to as NVL2-Δ8 - has not been described in the literature to date. However, in the annotation of NVL2 by ENSEMBLE annotation, a hypothetical splice variant with exon 8 exon skipping that has not been experimentally confirmed is predicted (NVL-208). Based on the sequence of NVL-201, exon skipping of exon 8 would result in a reading frame shift and a premature stop codon, thereby making NVL2-Δ8 a bona fide target of nonsense mediated decay (NMD) (Kervestin & Jacobson, 2012).This is consistent with the reduced stability of NVL2 transcripts in CMTR1-KO cells observed in RNA-seq (see Table 3).

**Table 3:**
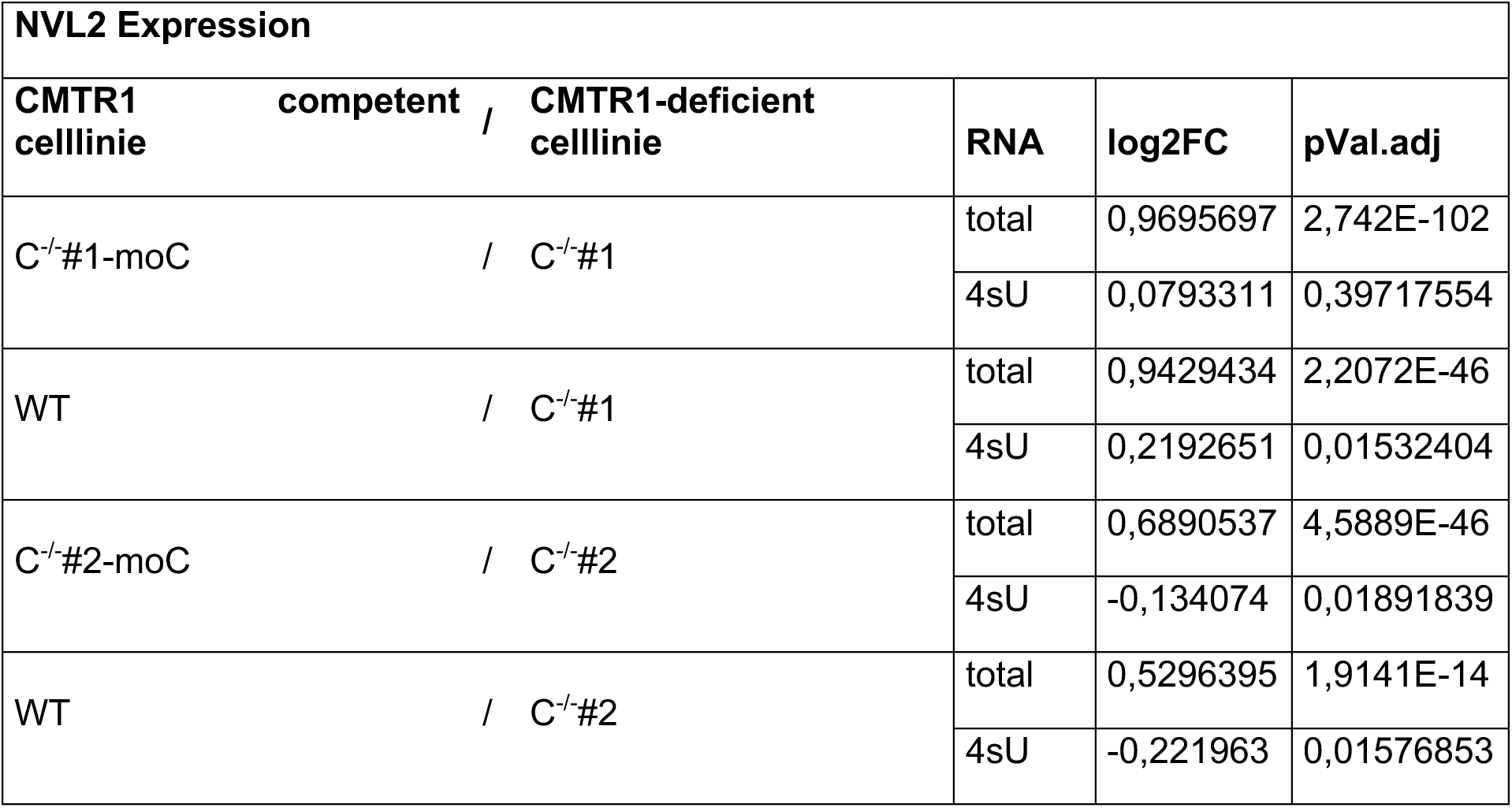
Differential expression of NVL2 between different CMTR1-competent and - deficient cell lines. RNA-seq data of differential transcript expression of NVL2 between the indicated comparison pairs (see Table 2).

NVL2 exon skipping was confirmed by qPCR and semiquantitative RT-PCR (sqRT-PCR) in RNA from untreated cells. Primer pairs with amplicons spanning either exon 8 and 9 or exon 9 and 10 were used for qPCR (Fig. 6D,E). While transcript levels of correctly spliced (i.e. exon 8 containing) NVL2 were significantly and strongly reduced in CMTR1^-/-^ cells compared with WT cells (Fig. 6D), total NVL2 transcript levels were only moderately reduced (Fig. 6E). SqRT-PCR with primers in exon 6 and 10 were used to contemporaneously detect correctly spliced NVL2 and NVL2-Δ8 and distinguished them based on different amplicon sizes. This demonstrated a reduction of correctly spliced NVL2 and the increased occurrence of NVL2-Δ8 in CMTR1-deficient cells (Fig. 6F,G).

### NVL2 expression depends on a catalytically active CMTR1

NVL2 is localized in the nucleolus and plays an important role in ribosome biogenesis as a member of the AAA-ATPase gene family (Kressler et al., 2012). NVL2 is known to interact with the nucleolar exosome to regulate the processing of rRNA and snoRNA (Hiraishi et al., 2018; LaCava et al., 2005; Yoshimatsu et al., 2015)). NVL2 function was investigated by Nahagama et al. using catalytically inactive NVL2 mutants (Nagahama et al., 2004). Expression of catalytically inactive NVL2 was observed to result in defects in 60S ribosomal subunit maturation and slowed processing of 47S pre-rRNA into 18S, 28S, and 5S rRNA (Yoshikatsu et al., 2015). It was shown by Hiraishi et al. that catalytically inactive NVL2 (NVL2-EQ) inhibited cleavage of 47S RNA at position (ITS1), which altered the kinetics of processing and resulted in altered intermediates (Hiraishi et al., 2018). Therefore, we analyzed rRNA maturation by Northern blot in CMTR1-competent and CMTR1-deficient cells. However, no difference in rRNA maturation between the tested cells could be detected (Suppl. Fig. 5). This finding suggests that CMTR1-deficient HEK293 cells do not exhibit a defect in rRNA processing despite greatly reduced endogenous NVL2 protein levels under normal growth conditions.

CMTR1 is a multidomain protein and dynamically interacts with other proteins (Inesta-Vaquera et al., 2018; Simabuco et al., 2019; Toczydlowska-Socha et al., 2018). Thus, loss of CMTR1 as a binding partner could disrupt interaction complexes and affect cellular processes independent of N1’2O-methyltransferase function. To analyze the impact of CMTR1 enzyme function and therefore N1’2O-methylation on the transcript levels of target genes, we transiently overexpressed wildtype (CMTR1-WT) and catalytically inactive (CMTR1-KA, amino acid exchange K239A (Belanger et al., 2010)) in the cells. Since NVL2 is involved in the ribosome biogenesis pathway, lack of NVL2 in CMTR1^-/-^ cells might be indirectly causing the observed reduction in transcript levels of f.e. RPs in CMTR1^-/-^ cells. To control for this, we also included conditions transiently overexpressing wildtype NVL2 (NVL2-WT) and catalytically inactive NVL2 (NVL-EQ, amino acid exchange E365Q and E682Q (Hiraishi et al., 2018)) in the experiment.

CMTR1-WT, CMTR1-KA, NVL2-WT and NVL2-EQ were overexpressed in WT, C^-/-^#1 and C^-/-^ #2 cells for 48h. After RNA extraction, transcript levels of NVL2, RPS12, PLCH1 and SNORA24 were determined by qPCR (Fig. 7). The overexpression of CMTR1-WT, but not other transgenes, was sufficient to significantly and strongly enhance the expression of correctly spliced endogenous NVL2.

**Figure 7:**
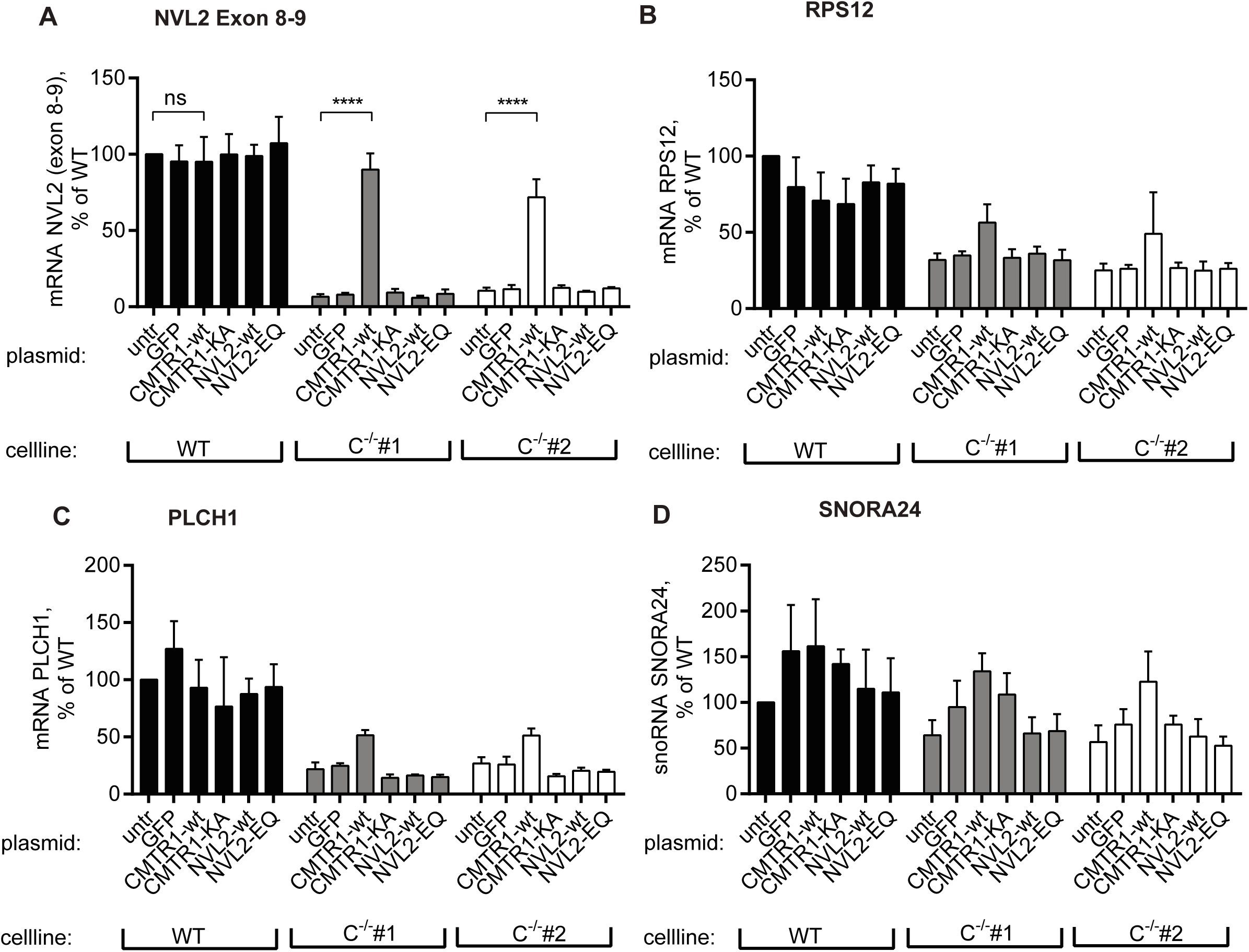
Effect of transgene overexpression on transcript levels in CMTR1^-/-^ cells. The cell lines WT, C^-/-^#1 and C^-/-^#2 were transfected with expression plasmids coding for the indicated transgenes and analyzed after 48 h by qPCR. Transcript levels of NVL2 exon 8-9 (A), RPS12 (B), PLCH1 (C) and SNORA24 (D) were normalized to GAPDH and are shown as % of untreated (untr) WT.

The other transcripts tested (RPS12, PLCH1, SNORA24) were identified in this work as downregulated in CMTR1-deficient cells (Suppl. Fig. 4). These transcript showed a slight, not significant increase in transcript levels upon CMTR1-WT expression, but not upon NVL2-WT or CMTR1-KA expression (7B-D). In addition, we could not determine an inhibitory effect upon expression of the catalytically inactive NVL2-EQ, which has been previously described to inhibit endogenous NVL2 in a dominant negative fashion (Hiraishi et al., 2018; Nagahama et al., 2004), indicating that NVL2 might not have the described biological function in our cells or is redundant in our cells.

Taken together, we show that the correct splicing pattern of NVL2 can be rescued by transient overexpression of catalytically active CMTR1, demonstrating that cap1-2’O-methylation is involved in splicing regulation of NVL2. In contrast, the biological consequence of the reduced NVL2 expression in CMTR1-deficient cells remains elusive.

## DISCUSSION

The cap0 modification of endogenous mRNA is strictly required for transcription, maturation, and translation in all eukaryotic cells. Multicellular organisms (metazoans) further introduce N1-2’O-methylation to form cap1 by CMTR1 (Belanger et al., 2010) So far, the hypothesis, that N1-2’O-methylation serves as a marker of endogenous mRNA for self versus non-self discrimination, is mostly based on the study of in vitro generated cap0/cap1 RNA and its differential recognition by RIG-I and IFIT1, and the fact, that N1-2’O-methyltransferase deficient viruses induce stronger RIG-I activation or are restricted by IFIT1 activation (Abbas et al., 2013; Daffis et al., 2010; Devarkar et al., 2016; Schuberth-Wagner et al., 2015; Zust et al., 2011).

Here, we use CMTR1^-/-^cell lines to characterize the biological function of the N1-2’O-methylation on endogenous mRNA. CMTR1^-/-^ cells have been described before, but have not been extensively characterized (Inesta-Vaquera & Cowling, 2017; Lee et al., 2020). While CMTR1^-/-^ guarantees the complete abrogation of endogenous N1 2‘O-methylation, it is limited to cell lines that survive long term loss of CMTR1^-/-^: Analysis of genome essentiality in human cells classified CMTR1 as an essential gene (Gurumayum et al., 2020). However, using artificial reporter cap0 or cap1 mRNA a recent study showed that differences in translational activity were heavily dependent of the cell line used (Sikorski et al., 2020). Therefore, the essentiality of CMTR1 catalyzed N1-2’O methylation of endogenous mRNA might differ between cell lines and depend on differences in RIG-I or IFIT1 expression and incomplete removal of CMTR1. This may explain the contradicting results about immune activation by endogenous RNA after RNAi or polyclonal KD of CMTR1: Li et al as well as our own previous work showed that CMTR1 depletion leads to a weak induction of IFN-beta transcripts, while the study of Williams et al. and this work show no effect on IFN transcription in untreated cells ((B. Li et al., 2020; Schuberth-Wagner et al., 2015; Williams et al., 2020). The low basal RIG-I expression in HEK293T cells which we used here is presumably not sufficient to sense endogenous RIG-I ligands in CMTR1^-/-^ cells and might have enabled the generation of viable CMTR1^-/-^ HEK293T cells and simplified the dissection of IFN-dependent and IFN-independent effects.

In this work, IFN-dependent effects on CMTR1^-/-^ cells were found to be largely mediated by IFIT1. IFIT1 restricts cap0 RNA from translation (Bartok & Hartmann, 2020; Daffis et al., 2010; Kumar et al., 2014). While IFIT1 shows no basal expression, it is highly induced after IFN-I treatment. Using cap0-or cap1 reporter mRNA, we showed that cap0 mRNA translation is equally reduced by IFN treatment depending on IFIT1 in WT and CMTR1^-/-^ cells, confirming that CMTR1^-/-^ cells have not aquired changes that enhance cap0 based translation.

CMTR1^-/-^ cells, but not CMTR1^-/-^ IFIT1^-/-^ or WT cells showed a dose-dependent reduction of global translational and viability after treatment with IFN-I.

Taken together, our results highlights the indispensability of CMTR1/cap1 for protecting endogenous mRNA against IFIT1 attack during IFN mediated antiviral responses.

This is in agreement with the findings of a recent study: Williams et al. analyzed 5 selected ISG after IFN-I treatment and observed a mildly reduced (two to threefold) induction of ISG15, MX1, IFITM1 protein after knockdown (KD) of CMTR1 in Huh7 cells. This effect was rescued by additional KD of IFIT1. Since only the protein level of a subset of individual ISGs were affected by CMTR1 knockdown, the authors propose that transcripts may be differently affected by IFIT1 function, possibly based on their 5’ secondary structure (Williams et al., 2020).

In contrary to Williams et al, we highlight the IFIT1-mediated inhibition of global translational activity in CMTR1 deficient cells, but it is indeed very likely that IFIT1-dependent translation inhibition will not affect all transcripts equally: IFIT1 binding was reported to be affected by further, less frequent cap methylations (N2 2’O, m6Am) or 5’ RNA secondary structures (ZITAT). Nevertheless. our data demonstrate that lack of CMTR1/cap1 generation is sufficient to cause an IFIT1 mediated global translational shut-down.,

While RIG-I downstream signaling and RIG-I, IFIT1, IFN-I and chemokine mRNA induction after RIG-I stimulation was intact in CMTR1^-/-^ cells, their translation was strongly reduced. This effect was reverted by depletion of IFIT1 (CMTR1^-/-^ IFIT1^-/-^ cells) and demonstrates that RIG-I dependent IFN-I production and the induction of adequate antiviral responses are strongly compromised in CMTR1 deficient cells.

This hypothesis is in agreement with results from Williams et al. who observed that Zika and Dengue virus replication is increased in cells with repressed CMTR1 expression (Williams et al., 2020).

Apart from the role within the antiviral immune response mediated by IFIT1 and possibly RIG-I, little is known about the biological significance of CMTR1 and N1-2’O-methylation of endogenous RNA. A couple of studies using RNAi of CMTR1 did not observe changes in global translation activity (Belanger et al., 2010) or nuclear export, RNA stability and polysome association of selected transcripts (Williams et al., 2020). Data from Inesta-Vaquera et al. show slowed cell growth after CMTR1 knock-down (Inesta-Vaquera et al., 2018). Negative data from knock-down studies are not easy to interpret since incomplete depletion of CMTR1 might hide significant functions.

We found the phenotype of CMTR1^-/-^ to be physiologically complex: CMTR1^-/-^ cells exhibited decreased cell size and slowed growth as well as altered expression of components of the translational machinery. The underlying mechanisms remain elusive. Based on the association of CMTR1 to the RNA Pol II CTD, we suspected that the loss of N1-2’O-methylation influences the transcriptome.

While the loss of CMTR1 potentially affects gene expression of all RNA Pol II transcripts, we found that gene global transcript stability and splicing were in fact not influenced by loss of CMTR1. However, we observed a significant reduction of RP mRNAs in CMTR1 deficient cells. Closer investigation revealed that apart from RP mRNAs other transcripts that posses attributes commonly associated with RP mRNA were also affected: Transcripts from non-ribosomal 5’TOP RNAs and SNHGs.

Most 5’TOP genes code for RPs or other proteins that are directly involved in the cellular translational machinery. 5’TOP transcripts are structurally defined by the eponymous 5’ terminal oligopyrimidin tract (5’TOP) of 5-14 pyrimidins. Functionally, 5’TOP transcripts are translationally regulated via the mTOR pathway (Meyuhas & Kahan, 2015). Unfavorable growth condition or inhibiton of mTOR leads to translational repression that especially affects 5’TOP transcripts. It is noteworthy that 5’TOP genes use a conserved transcription start site (TSS) within their core promoter, thereby initiating with C as the first nucleotide (N1), while the vast majority (86% in HEK293T) of RNA Pol II transcripts are described to initiate with A or G (Philippe et al., 2020). While eIF4E has been shown to have lower affinity for N1=C transcripts, thereby possibly contributing to the translational inhibition of 5’TOP transcripts via mTOR, it is unclear whether the identity of the N1 plays a role in N1 2’O methylation by CMTR1 and mRNA maturation in general. If lack of CMTR1 specifically affects transcription of N1=C transcripts, these effects would unlikely be detected by changes in RNA stability as measured here, since transcription of incompletely capped transcripts is aborted at the promotor proximal pausing step. Surprisingly, a recently published study shows that a large proportion of genes classified as 5’TOP can use alternative TSSs, thereby deviating from N1=C (Nepal et al., 2020). While the biological significance of this observation is still largely unclear, it has been shown that TSS usage can vary under experimental conditions and that effective snoRNA maturation relies on 5’TOP-specific TSS usage of the SNHG transcript (de Turris et al., 2004; Nepal et al., 2020).

It would be conceivable that changes in the cap methylation status in CMTR1 deficient cells exerts differential effects on 5’TOP RNA processing depending on the initiation used, leading to the reduced expression of 5’TOP RNA, SNHG RNA and snoRNA in CMTR1-deficient cells. Furthermore, TSS usage could be altered as a consequence. To prove this, the TSSs actually used would have to be determined experimentally in CMTR1-/- cells.

We found no evidence of changes in mTOR regulation in CMTR1 deficient cells (data not shown). In contrast to translational regulation of 5’TOP RNAs, regulation of 5’TOP RNAs which are usually used as house keeping genes by transcript level as found in this work are uncommon. Interestingly, the lack of CMTR1 phenocopies the inactivation of mTOR regulated effector proteins in regard of the cell size: Inactivation of 4E-BP and S6 kinase has been described to result in reduced cell size (Fingar et al., 2002; Pollizzi et al., 2015; Ruvinsky et al., 2005). In a recent study knockdown of CMTR1 was observed to reduce histone and ribosomal protein gene expression in embryonic stem cells (S. Liang et al., 2022) and ChIP experiments revealed less RNA-Polymerase II binding to the corresponding promoters suggesting that CMTR1 acts as a transcription factor. However, the requirement of enzymatic activity was not tested and the effect was dependent on ESC differentiation. Thus an indirect effect of CMTR1 on expression/splicing of another transcription factor cannot be excluded. Proteome analysis revealed, that despite the observed regulation of RP mRNA, RP protein expression appears to be reduced but this effect is not significant. However, we found the ribosome biogenesis factor NVL2 to be strongly downregulated on protein level.

While we could not determine the influence of CMTR1 on global splicing efficiency, we found that the decreased NVL2 protein levels in CMR1^-/-^ cells to be caused by a highly specific and previously undescribed exon skipping event of exon 8 of the NVL2 gene (NVL2-Δ8). The affected splice sites in the NVL2 gene and their genomic context were analyzed with the software Splice Port (Dogan et al., 2007), but did not show any obvious special features (data not shown). It could not be elucidated how the CMTR1 status of the cell controls the alternative incorporation of this exon.

NVL2 has been reported to be implicated in rRNA processing (Hiraishi et al., 2018; Nagahama et al., 2004; Yoshikatsu et al., 2015) and to be an essential gene (Gurumayum et al., 2020). It is tempting to interpret the totality of CMTR1-dependent effects - reduced cell size, slowed proliferation and reduced expression of components of the translational machinery - as compensation by the cell for reduced efficiency of ribosome biogenesis. Based on the function described in the literature, NVL2 is a promising candidate that could account for these effects. However, although NVL2 protein expression was drastically reduced in CMTR1-deficient cells, we observed no change in rRNA processing. Apparently, low NVL2 levels are sufficient to maintain ribosome biogenesis at normal growth conditions. While transient overexpression of CMTR1 in CMTR1-deficient cells restored the expression of correctly spliced NVL2 as well as partially rescuing the transcript levels of RPs, SNHGs and snoRNA, overexpression of NVL2 showed no comparable effect. Thus, it is unlikely that the reduced protein expression of NVL2 in CMTR1-deficient cells has a direct impact on the altered transcript levels of RPs, SNHGs and snoRNA.

It should be noted, that the usage of CMTR1^-/-^ deficient cells in this work compared to transient CMTR1 inactivation ensures the lack of endogenous N1-2’O-methylation, thereby enabling the discovery of new functions of N1-2’O-methylation of endogenous mRNA as demonstrated in this work. On the other hand, the use of stable CMTR1^-/-^ may lead to an underappreciation of the effects of N1-2’O-methylation, since potential lethal effects of CMTR1^-/-^ cells would either impair KO clone generation or have to be compensated for by the cell. This is especially relevant for our study, since both CMTR1^-/-^ and our main candidate NVL2 are involved in fundamental cellular processes of transcription, translation and ribosome biogenesis and are described to be essential genes. We address these issues by including appropriate controls.

While the basics of transcription and RNA maturation have long been known, their precise kinetics and regulation remain a highly active field of research. An understanding of how complex, tight and reciprocal transcription and RNA processing steps such as capping and splicing are coupled is only slowly emerging (Kachaev et al., 2020; Tellier et al., 2020). The insights gained in this work provide new insights into the complex role of N1-2’O-methylation in gene expression and splicing in higher eukaryotes, while the underlying mechanism remain to be elucidated.

## MATERIAL AND METHODS

### Cell lines

HEK293T were cultivated in DMEM (10%FCS, 100 U/ml penicilin, 100 mg/ml streptomycin)at 37 °C.

HEK-Blue™IFN-α/βwere bought from InvivoGen and cultivated in DMEM (10%FCS, 100 U/ml penicilin, 100 mg/ml streptomycin)at 37 °C

### Cell culture

HEK293T cells and the generated in this work celllines based on HEK293T were cultured in cellculture dishes and harvested by trypsination, counted and a desired cellnumber was plated in appropiate cellculture dishes, depending on the experimental setup. For analysis of metabolic activity, luciferase activity or cytokine production 96well plates were used, qPCR or sq-RNA analysis was performed from cells plated in 12 well plates, westernblot analysis or colony formation assays were performed in 6 well plates. The cellnumber used is indicated in the Fig. legends.

Transfections with DNA or RNA were performed using Lipofectamine 2000 (LF2000, Thermo Fisher Scientific, Darmstadt, Germany) according to manufacture protocol with a maximal ratio of 500 ng nucleic acid to 1ul Lipofectamin. In brief, for the transfection of a single well of a 96 well plate 25ul OPTImem with 0,2ul Lipofectamin was preincubated for 5min at RT and subsequently mixed with up to 100ug nucleic acid diluted 25ul Opti-MEM. The transfection mix was incubated for 20 min before it was added to the cells. This transfection protocol was scaled up for transfection of other cell culture palte formats.

### Generation of gene-edited and reconstituted cell lines

20*10^5^ HEK293T cells were plated in 96 well plates. After 24h the cells were transfected with LF2000 with 100 ng of a CAS9-gRNA plasmids, targeting CMTR1 (5’GGGTCGGAAGGACATCGTTG(AGG)3’) or IFIT1 (5’AGGCATTTCATCGTCATCAA(TGG)3’). 24h after transfection, the cells were trypsinized and seeded in limited dilution to obtain single-cell clones. After expansion knockout (KO) was confirmed by immunoblot or Sanger sequencing. For stable reconstitution of CMTR1 expression, murine CMTR1 cDNA was cloned into the retroviral expression vekor pRp-puro and retroviral particles were generated (Stutz et al., 2013). Target cells were transduced with retroviral particles and stable transgene expressing cell lines by puromycin selection (2ug/ml final in culture medium).

### RIG-I stimulation and quantification of cytokines

We used triphosphorylated, doublestranded RNA Ligand IVT4 for stimulation of RIG-I signalling (Goldeck et al., 2014). RIG-I mediated cytokine induction was measured 24h after stimulation. IP-10 levels were analyzed with commercial ELISA assay kit (Thermo Fisher Scientific) according to the manufacturer’s protocol. Type I Interferon (IFN I) levels were determined using the reporter cellline HEK-Blue™IFN-α/β (INVIVOGEN) according to manufacturer’s protocol. The concentration of cytokines in the sample was determined by a standard curve generated with recombinant IP-10 (Thermo Fisher Scientific) or rekombinant IFN-α (IFN-a2a, Milteny).

To measure the RIG-I activation with the IFN reporter system, the cells were co-transfected with the indicated amount of IVT4, 15 ng Gaussia luciferase (GLuc) reporter plasmid (pGL3-IFNB1-GLuc) and 15 ng control Firefly Luciferase expression plasmid for normalization (pLenti-Ef1α-FLuc). After 24h the luciferase assay was performed in white microplates and luminescence was measured with the EnVision 2104 Multilable plate reader (Perkin Elmer). GLuc production was measured as a surrogate for IFN-β promoter activation by adding 25 µl coelenterazine-solution (1 µg/ml in H_2_O) was to 25 µl supernatant.reader (Perkin Elmer, Waltham, USA) and luminescence was measured. To measure FLuc as a control, the supernatant was aspirated and 40 µl 1 x SAP-solution (0.02 % Saponin, 50 mM tris HCL pH 7.8, 15 mM MgSO 4, 4 mM EGTA, 10 % Glycerol) was added to the cells and then they were lysed for 20 min at RT on a shaker. Afterwards, 25 µl of lysed cells in SAP were added to 25 µl Firefly luciferase substrate (Luciferin) and luminescence was measued.

### Colony formation assay

Colony formation assay was performed in 6 wells plates that had been coated by poly-L-ornithine (1 mg/ml in H2O) incubation for 1h at 37°C. The coating solution was aspirated and the plates were washed two times with PBS. 5×10^5^/well HEK293T cells were plated and immediately treated with 10.000 U/ml IFN-α or left untreated. After 10 d, the cell culture medium aspirated and the cells were washed with PBS and subsequentially fixed for 30 min at room temperature with 1 ml 4% formaldehyde in PBS. The fixative solution was removed and the cells incubated with 1 ml 0.05% crystal violet solution for 1 h at room temperature. The staining solution was removed and the plates were washed with water by repeated washing steps until a good contrast between colonies and background was achieved. The water was then completely removed and the plates dried at room temperature. The plates were scanned with Odyssey SA at 700 nm. By subsequently solubilising the dye, the relative amount of bound dye well can be quantified photometrically (Franken et al., 2006). For this purpose, the plates were incubated with 1 ml/well 10% SDS for 24 h at 37°C in a cell culture incubator. 200 μl was transferred to a 96-well microtitre plate and the absorbance was measured at OD584nm in the Epoch plate spectrometer.

### Measurement of cell growth kinetics and cell viability by MTT Assay

MTT assay is based on the reduction of thiazolyl blue (MTT) to purple formazan crystals in intact cells (Berridge & Tan, 1993). The turnover of MTT depends on the metabolic activity of the individual cells and the total number of cells in the well and can be used as an indicator of cell viability, cytotoxicity or proliferation, depending on the application. Cells were cultured in 96-well cell culture plates in 100 μl cell culture medium.

To determine effect on cellviability, cells were treated with a titration of IFN-α or inhibitors and analyzed at the indicated end point. 20 μl MTT substrate (5 mg/ml MTT in PBS) was added to the culture medium and incubated at in the cellculture incubator until a distinct staining of the untreated cells was visible (30 to 90 min). The reaction was then stopped by adding 100 μl 10% SDS to solubilize the formazan crystals and photospectrometrically analyzed at OD 584nm in the Epoch plate spectrometer.To measure cell proliferation as a function of time in culture, untreated cells were incubated with 20 μl MTT substrate for exactly 1 h at the indicated time points and then lysed with 100 μl 10% SDS and analyzed as described above.

### FACS based analysis

FACS experiments were performed with LSR II (BD) and analyzed with the FlowJo 8.7. Prior to measurement, cells were washed once with PBS and taken up in FACS buffer (2 mM EDTA and 2% FCS in PBS). For the analysis, the population of intact single cells was selected in the FSC and SSC, cell fragments were excluded. Cell cycle analysis requires fixation and subsequent staining with propidium iodide (PI), an intercalating DNA-binding fluorescent dye, to determine the cell cycle phases of cells due to the DNA content of the cells. 1-5×105 cells were washed two times with FACS buffer and resuspended in 100 μl MACS buffer. The cells were fixed by addition of 1 ml ice-cold 100% ethanol, mixed by inverting and incubated for 16 h at 4°C. The ethanol fixed cells were pelleted by centrifugation for 5 min at 800 RCF and then resuspended 1 ml MACS buffer. Afterwards, the cells were pelleted (5min 500 RCF) and washed with MACS buffer two times. The pellet was resuspended in 100 μl MACS buffer containing 100µg/ml RNAse A and 10 µg/ml PI and incubated for 30 min in the dark. Subsequently, the cells were analyzed by FACS as described above, but including measurement of the fluorescence intensity of the PI channel.

### RNA Synthesis

The RIG-I Ligand IVT4 was prepared as described before (Goldeck et al., 2014).

The generation of triphosphoryted RNA Oligonukleotides with subsequent enzymatic capping were performed as described before(Schuberth-Wagner et al., 2015).

Reagents to prepare Cap0- and Cap1-modified Renilla Luciferase (RLuc) mRNA using the CleanCap technology (Trilink)were provided as part of a cooperation with Trilink Bioscience. The CleanCap technology is based on a Cap0- or Cap1-modified dinucleotide (m7GpppAG or m7GpppAmG), which can be co-transcriptionally integrated into in vitro transcripts (IVT). The cap0 and Cap1 modified RLuc mRNA were prepared according to the manufacturer’s protocol.

### RNA based analysis

For PCR based analysis, cellular RNA was prepared was extracted using Zymo III columns (Zymo, Freiburg, Germany). Reverse transcription was performed using random hexamer primers (IDT) and RevertAid Reverse Transcriptase (Thermo Fisher Scientific). qPCRs was performed using 5x EvaGreen QPCR-Mix II (ROX) (Bio-Budget, Krefeld, Germany). Primers were validated with melting curve analysis and tested for efficiency using cDNA dilution.

Semiquantitative reverse transcriptase PCR (sqRT-PCR) was used to discriminate between alternative splicing products of NVL2 by amplicon size. The sqRT-PCR uses cDNA, prepared as described above, as a template. sqRT-PCR was performed in 25 μl reaction volume, consiting of 2.5 μl 10x DreamTaq Green Buffer, 21.2 μl water, 0.2 μl each of forward (fwd) and reverse (rev) primers (10 mM), 0.2 μl DreamTaq DNA polymerase and 0.5 μl cDNA. The PCR reaction was performed in the thermal cycler with 2 min 95°C (initial denaturation), 30x amplification cycles consisting of 15 sec 95°C, 15 sec 59°C and 45 sec 72°C. After a further 10 min at 72°C (final elongation), the reaction mix was cooled to 4°C and analyzed by standard agarose gel electrophoresis with 2% (w/v) Agarose in TAE Buffer with 1:20.000 SybrSafe (Thermo Fisher), followed by imaging with Odyssey Fc at 600nm.

For other analysis, cellular RNA was prepared TRIzol-based RNA extractions (Thermo Fisher Scientific) according to manufacturer’s protocol and solubilize in H_2_O. RNA concentration and quality was determined photospectrometrically.

Digestion of cellular RNA preperations with Terminator™5’Phosphate-Dependent Exonuclease (Epicentre) was perforemd to remove the abundant rRNA, thereby increasing the concentration of other RNA Species. Up to 10μg cellular RNA was digested with terminator nuclease according to the manufacturer’s instructions (30 min at 37°C), the reaction was terminated by adding 1 ml TRIzol purified as described above.

### MTase-Glo based assay for the determination of cap methylation

In order to determine the cap methylation status of RNA, an assay was developed in this work to detect cap0-modified RNA. For this purpose, the enzymatic activity of the commerically availablale recombinant vaccinia virus methyltransferase mRNA 2’O-methyltransferase (NEB), abbreviated hereagter as VMTR1, was used. This VMTR1 requires Cap0 RNA as a substrate and generates Cap1 (Barbosa & Moss, 1978), consuming the methyl donor S-adenosyl-methionine (SAM) and converting it to S-adenosyl-homocysteine (SAH). SAH can be quantified by the MTase-Glo assay (Promega). Quantification is based on the conversion of SAH to adenosine diphosphate (ADP), which generates a luminescent signal (Hsiao et al., 2016).

10 μl of sample RNA dilution for the reaction and for the mock reaction were prepared in reaction tubes(2,000 ng TRIzol-purified RNA from cells, 500 ng terminator nuclease-treated RNA from cells, 25 ng capped RNA oligonucleotides). 10 μl enzyme mastermix (5 μl MTase Glo 4x Reaction Buffer, 2 μl SAM (100 μM), 0.2 μl VMTr1 and 2.8 μl water) or mock-mastermix, lacking VMTR1, were added to the RNA dilution and and incubated for 1h at 37°C, then 25 μl MTa-Glo Detection Solution (Promega) was added and incubated for a further 30 min at room temperature. Finally, 45 μl of the reaction were transferred to a LumiTrack-200 96-well plate and the luminescence read in the EnVision 2104 Multilabel reader. The relative SAM consumption was determined as the quotient of the luminescence between a reaction with methyltransferase and a mock reaction (without enzyme).

### Immunoblot

Immunoblot was measured to study protein levels. The cells were cultured in 6 well dishes and lysed in 100 µl 1 x Laemmli-buffer. (120 mM Tris pH 6.8, 4 % (w/v) SDS, 20 % (v/v) glycerol, 20 mM DTT and Orange G). Samples were sonificated and heated (95 °C, 5 min) prior to loading.

After transfer, the membrane was stained with Ponceau-S solution (0,1% (w/v) in 5% acetic acid) to quantify the amount of loaded protein. After destaining with H_2_O, immunoblot was performed with the primary and secondary antibodies are listed in the resources table. Blots were recorded on an Odyssey FC Dual-Mode.

Imaging system (LI-COR Biosciences, Bad Homburg, Germany).

#### Northern blotting

Total RNA was extracted using Trizol according to the manufacturers protocol. 4 µg of total RNA was resolved on a 1% agarose/0.4 M formaldehyde gel using the tricine/triethanolamine buffer system (30 mM tricine, 30 mM triethanolamine). The RNA was blotted on an uncharged nylon membrane (Carl Roth, ROTI®Nylon 0.2; Art. No. AE50.1) by capillary transfer overnight in 10x SSC buffer (1.5 M sodium chloride, 0.15 M sodium-citrate; pH 7.0). Following UV-crosslinking (0.12 J/cm2 each membrane side with Vilber Lourmat BLX-254), the blots were prehybridized for 1 h in Church buffer (0.5 M di-sodium hydrogen phosphate, 1 mM EDTA, 7% SDS, pH 7.2). Hybridization with 5’-32P-labeled oligonucleotides (labelled with PNK and γ-32P-ATP [800 Ci/mmol, 10 mCi/ml]) in Church buffer was carried out overnight at 40 °C. The oligonucleotide sequences were as follows: ITS1: (5′-CCTCGCCCTCCGGGCTCCGTTAATGATC-3′), ITS2: (5′-CTGCGAGGGAACCCCCAGCCGCGCA-3′), 5 ETS: (5′-CGACAGGTCGCCAGAGGACAGCGTGTCAGC-3′). All blots were washed 2x with Wash buffer 1 (2x SSC, 0.1% SDS) and 2x with Wash buffer 2 (0.2x SSC, 0.1% SDS) at 40°C for 15 min each. RNA signal from at least three distinct samples were detected with the Typhoon FLA 7000 (GE Healthcare) and were quantified in a semi-automated manner using the ImageQuant TL 1D software (version 8.1) with a rolling-ball background correction. Ethidium bromide stained 28S and 18S rRNA served as loading controls. The control condition was set to unity (wt), quantification results are shown as data points and mean.

### SUnSET Assay

The SUnSET (surface sensing of translation) assay allows quantification of the translational activity of cells (Goodman & Hornberger, 2013; Schmidt et al., 2009). The SUnSET assay is based on the incorporation of puromycin into nascent proteins, which can subsequently be detected and quantified in the immunoblot by an anti-puromycin antibody. Cells were spiked with puromycin (final concentration 1 μg/ml) for 20 min and incubated at 37°C for 20 min. Afterwards, the cells were lysed and processed by standard immunoblot methods. the incorporation of puromycin was measured using an anti-puromycin primary antibody.

### RNA-Seq

The transcriptome of the cellines WT, C^-/-^#1, C^-/-^ #2, C^-/-^#1+moC and C^-/-^#2+moC was analyzed by RNA Sequencing. In addition to total cellular RNA, newly transcribed RNA was prepared by metabolic labeling with 4sU (carbosynth) and analyzed.

Cells were seeded 2 d before 4sU labelling in 15 cm cell culture dishes. The number of cells used was adjusted according to the growth rate of each cell line to ensure a cell density between 50 and 75% at the start of 4sU labelling. A final concentration of 500 μM 4sU (carbosynth) was added to the cells and incubated for 1 h in the cell culture incubator at 37°C. The 4sU-containing medium was then completely removed and the cells lysed by additions of 7.5 ml TRIzol, then cellular RNA (total RNA) was purified. Preparation of metabolic 4sU labeled RNA (4sU RNA) out of the total RNA relies on sulfhydryl-specific biotinylation and Strepvidin based purification and was performed according to published protocols (Dölken et al., 2008; Rutkowski et al., 2015):

The 4sU RNA and total RNA were processed and sequenced (HiSeq 4000, 5PE100) by the sequencing service provider BGI for library production (BGI eukaryotic long non-coding RNA sequencing, lncRNA library). Sequencing quality was assessed with FastQC (http://www.bioinformatics.babraham.ac.uk/ projects/fastqc/) and sequencing adapters were trimmed using Cutadapt (Martin, 2011). RNA-seq reads were mapped against the human genome (hg38) and human rRNA sequences using ContextMap version 2.7.9 (Bonfert et al., 2015) (using BWA (H. Li & Durbin, 2009) as short read aligner and with default parameters). Number of read counts per gene and exon were determined from the mapped RNA-seq reads in a strand-specific manner using featureCounts (Liao et al., 2014) and gene annotations from from Ensembl version 91 (Zerbino et al., 2018). The differential gene expression analyzes were performed with DESeq2 (Love et al., 2014). Only transcripts that had at least 25 reads in both of the conditions to be compared were considered. P-values were adjusted for multiple testing using the method by Benjamini and Hochberg (Benjamini & Hochberg, 1995). To identify differential alternative splicing events, differential exon expression analysis was performed using DEXSeq (Anders et al., 2012). Analysis workflows were implemented and run using the Watchdog workflow management system (Kluge & Friedel, 2018).

The efficiency of the splicing response of individual cell lines and conditions was quantified by determining the global "splicing index". For this purpose, the reads spanning the splice site are first classified, based on their sequence information, into "exon-exon" reads (which indicate a completed splicing reaction of the RNA) and "exon-intron" reads (which indicate unprocessed splice sites in the RNA). The splicing index was calculated using the following formula:

Splicing index= (reads(exon exon))/(reads (exon exon)+reads (exon intron)).

The averaged splicing index of all splice sites of a gene resulted in the splicing index of the gene, of which the average was determined across the replicates. The global splicing index of the cell line in turn results from the mean value of the splicing index of all measured genes. For the analysis of transcript stability, the quotient of transcript expression in FPKM (fragments per kilobase of exon model per million reads mapped) between newly synthesised RNA (4sU) and total RNA (total) was calculated for all detected transcripts. The mean value of the replicates was calculated and the 4sU/total quotients of all detected transcripts were plotted against each other to compare the transcript stabilities between different cell lines. GO Enrichment was performed with the PANTHER web application (Overrepresentation Test, http://pantherdb.org).Volcanoplots were created using the program R (R version 3.6.1) with the graphical interface R Studio (version 1.0) and ggplot2.

### Proteome analysis

Explorative proteome analysis C^-/-^#2+moC and C^-/-^#2 were performed by mass spectrometry with label free quantification (LFQ) according published protocols. Samples were generated by plating 3×10^5^ cells in technical tetraplicates in 6-well plates. After 24 h the cells were washed with PBS 2 times and detached by pipetting. After pelleting by centrifugation at 500 rpm for 5 min the the cell pellet was frozen in liquid nitrogen. The analysis was performed by Alexey Stukalov (AG Pichlmair, Institute of Virology, TU Munich). The volcanoplot were generated using the program R (R version 3.6.1) with the graphical interface R Studio (version 1.0) and ggplot2.

## ACKNOWLEDGEMENTS

This study was funded by the Deutsche Forschungsgemeinschaft (DFG, German Research Foundation) under Germany’s Excellence Strategy – EXC2151 – 390873048 of which GH and MS are members. It was also supported by other grants of the DFG, including TRR237 (GH, MS), SFB670 (GH, MS), and DFG SCHL1930/1-2 (MS). MS received financial support from BONFOR (University of Bonn). This work is part of the PhD thesis of SW at the University of Bonn.

## AUTHOR CONTRIBUTIONS STATEMENT

SW, TH, CW, APir, VB, ME, SJ and MS performed the experiments. SW, TH, VB, ME, AP, LD, CF and MS conceived experiments. SW, VB, NG, CF, GH, AP, LD, AS, CF and MS analyzed and interpreted data and wrote the manuscript. All authors revised the manuscript.

---

**Supplementary Figure 1:**
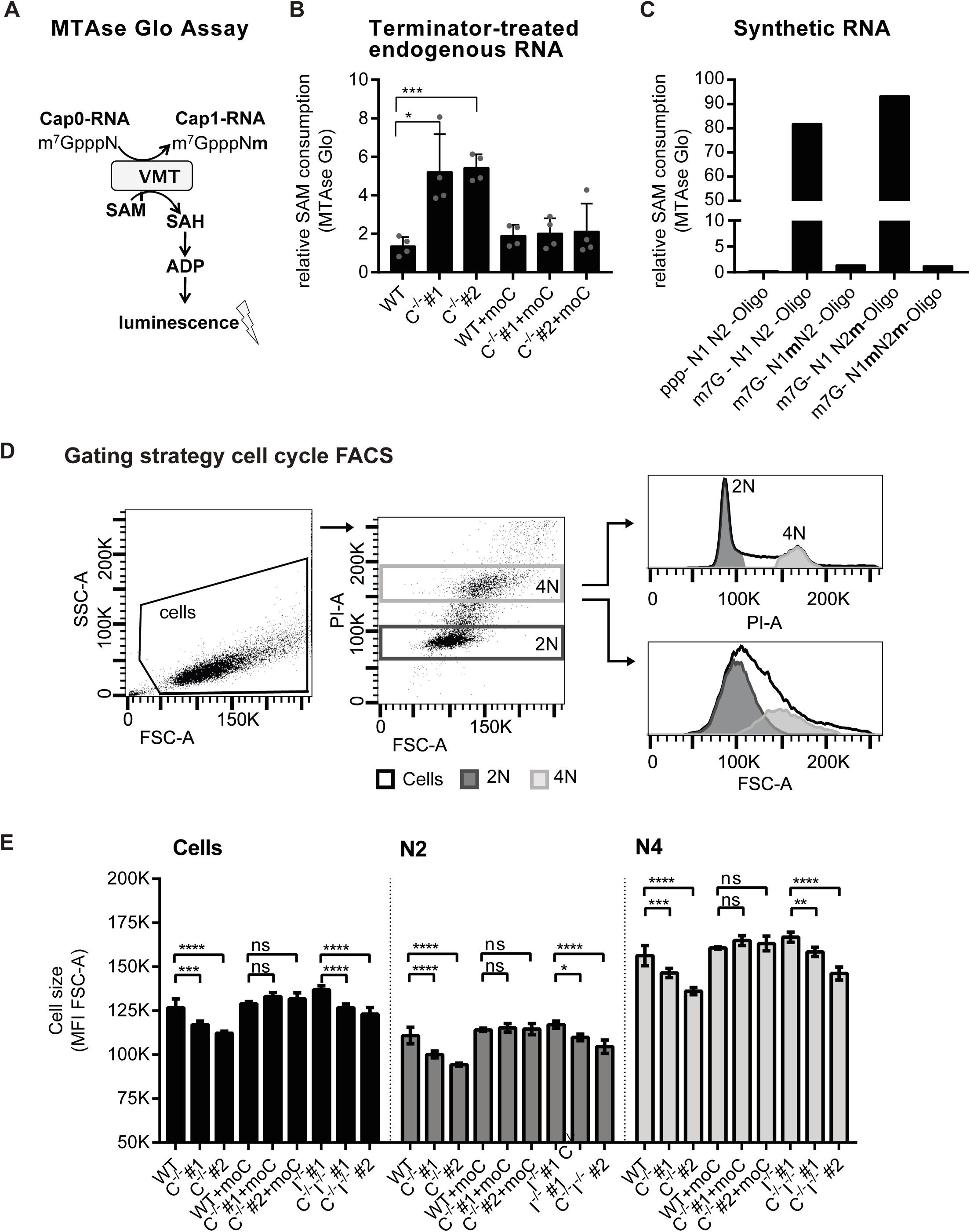
A-D) Detection of endogenous cap0-modified RNA with the MTAseGlo Assay. A) schematic depiction of the MTAse Glo Assay B-C) MTAse Glo assay was performed as described in Figure 1 and methods. B) Validation of the assay specifity for cap0-modified RNA species with modified RNA oligonucleotides (25ng) (representative result). C) MTAse Glo assay with Terminator nuclease treated cellular RNA (500ng). D/E) Flow cytometric analysis of cell cycle phases and cell size. Ethanol fixed, propidium iodide (PI) stained cells were analysed. D) Gating strategy shown for a representative experiment. The population "cells" was defined in the FSC/SSC, and within this population, populations 4N and 2N were defined based on PI staining. E) Statistical analysis of cell size in mean fuorescence intensity (MFI) of FSC-A per population (cells, 2N and 4N) (n=4, mean + SEM).

**Supplementary Figure 2:**
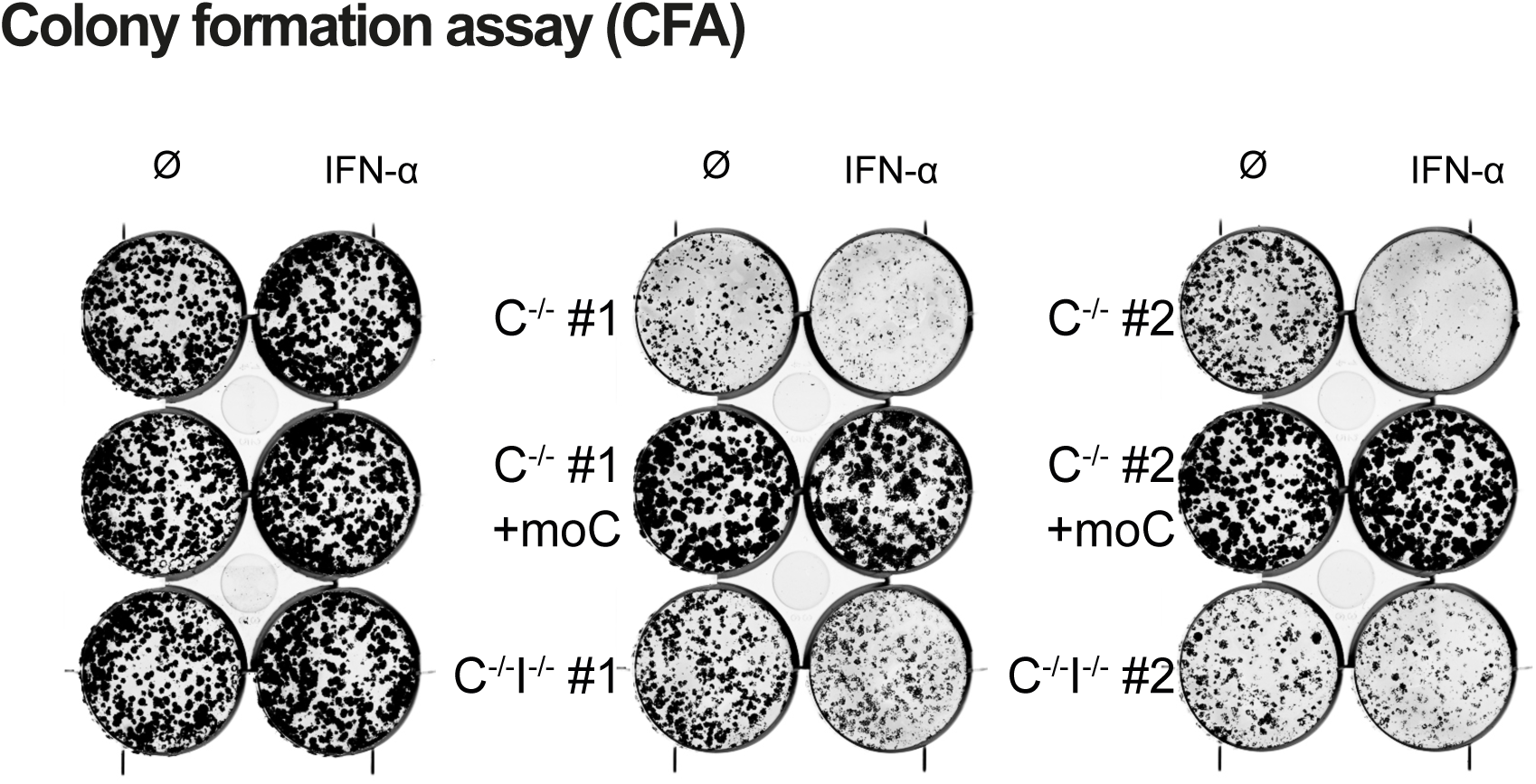
IFN-I/IFIT1 dependent and independent effects of CMTR1 on cell growth. A) Colony formation assay (CFA). 5×10^3^ cells/well were seeded in 6-well plates and left untreated (∅) or incubated with 10.000U/ml IFN-∝. After 10 days, crystal violet staining was performed and cell plates were scanned at 700nm. A representative result is shown.

**Supplementary Figure 3:**
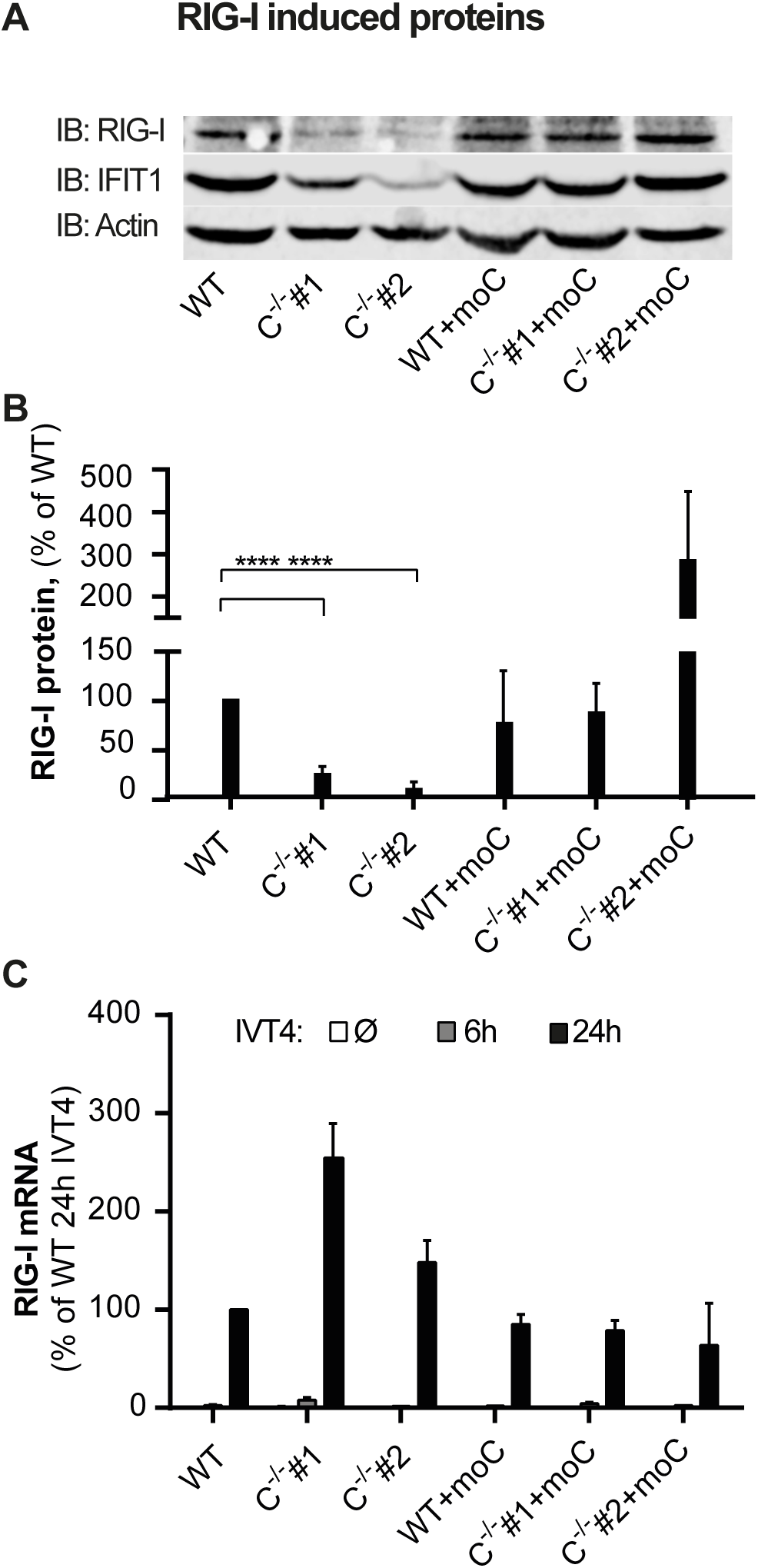
No effect of CMTR1 on signal transduction but translation down-stream of RIG-I. A/B) Indicated cells were stimulated for 24 h with IVT4 (0.5 µg/ml). A) Representative experiment, WB of RIG-I IFIT1 and actin. B) Quantification of the relative amount of R G-protein amount normalized to actin (n=6), mean + SEM. C) qPCR analysis of cells stimulated for 6h or 24h with IVT4 (0.5 µg/ml) or left untreated. Relative mRNA expression of R G-normalized to GAPDH (n=4, mean + SD).

**Supplementary Figure 4:**
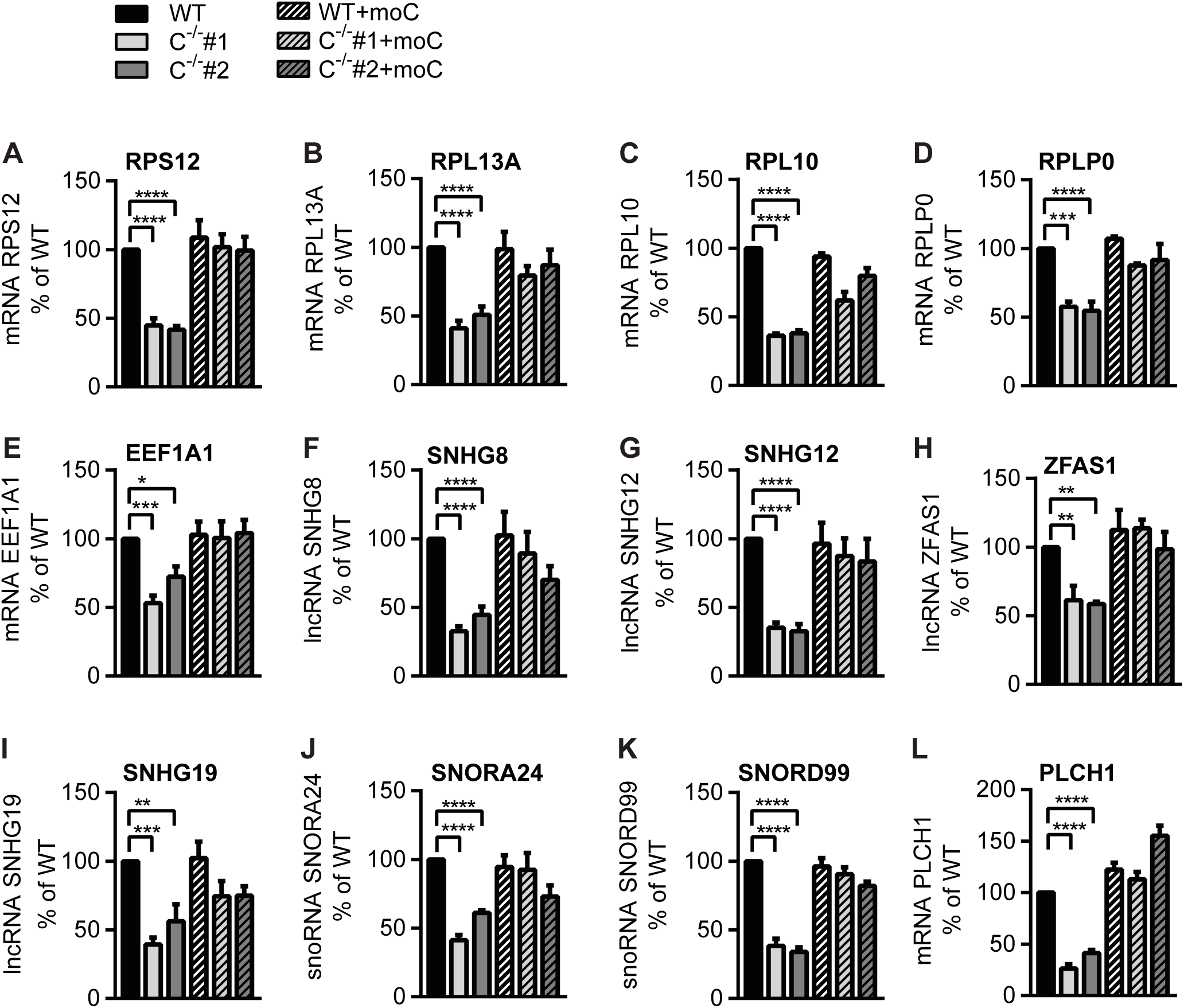
Validation of the differentially regulated genes in CMTR1^-/-^ cells. qPCR analysis of RNA from untreated cells. The relative transcript levels of RPS12 (A), RPL13A (B), RPL10 (C), RPLP0 (D), EEF1A (E), SNHG8 (F), SNHG12 (G), ZFAS1 (H), SNHG19 (I), SNORA24 (J), SNORD99 (K) and PLCH1 (L), normalized to GAPDH, in % of WT (n=4, mean + SEM).

**Supplementary Figure 5:**
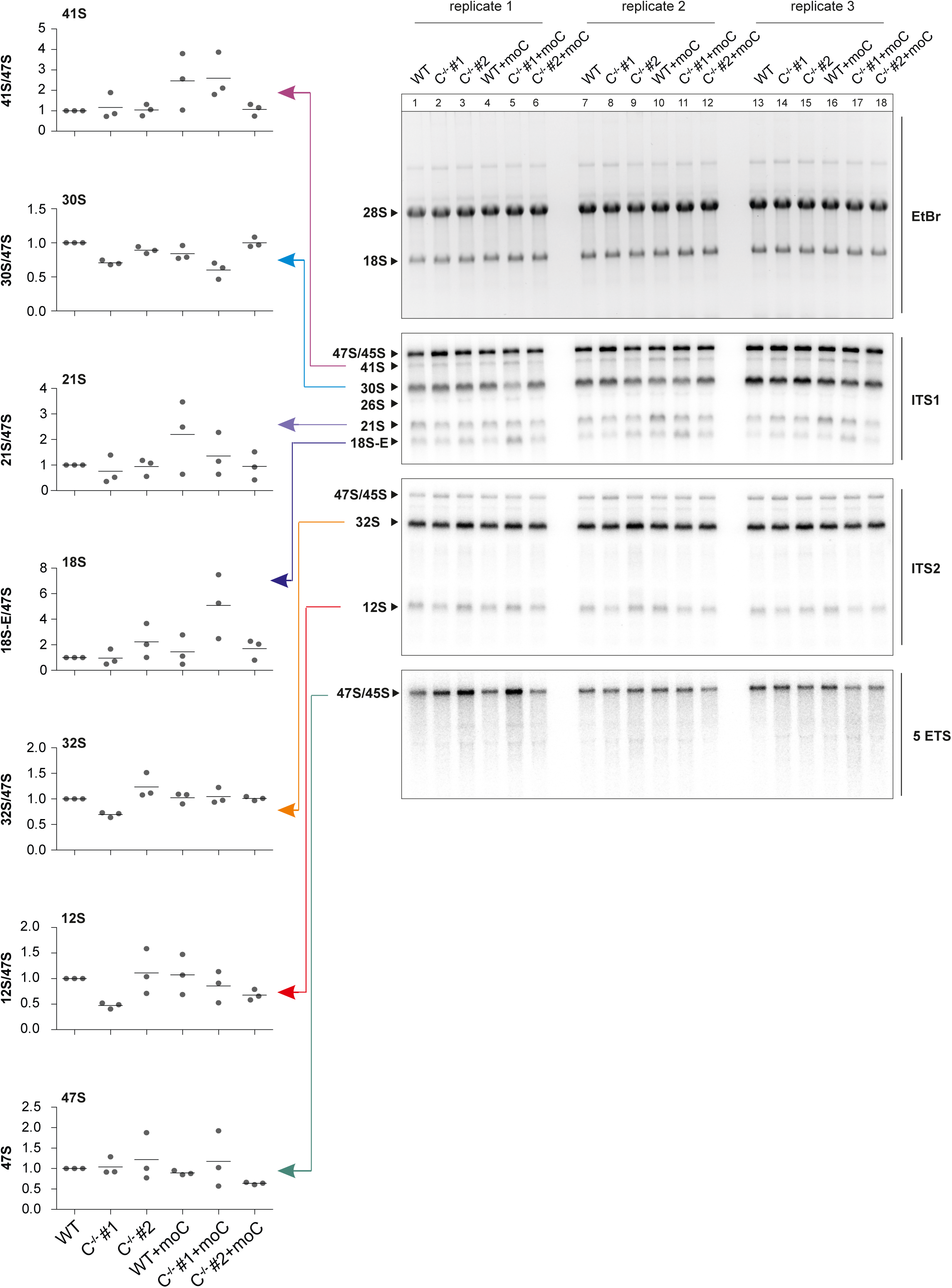
rRNA processing is not significantly impaired in CMTR1^-/-^ cells. Northern blots of total RNA obtained from indicated cell lines. The hybridization was performed with probes targeting the ITS1, ITS2 or 5’-ETS (5 ETS) region of ribosomal precursor RNAs. Ethidium bromide (EtBr) stained 28S and 18S rRNA served as loading controls. Quantification of the northern blots shows the amounts of the indicated precursor rRNAs normalized to the corresponding amount of 47/45S pre-rRNA. Data points and mean are shown (n=3).

